# Modeling social influence as a reinforcer reveals varying individual learning strategies and the group’s structure

**DOI:** 10.1101/2025.11.06.686867

**Authors:** Michał Lenarczyk, Bartosz Jura, Zofia Harda, Łukasz Szumiec, Magdalena Ziemiańska, Jan Rodriguez Parkitna, Daniel Krzysztof Wójcik

## Abstract

In a social context, knowledge can be gained through observation and imitation, but in a numerous group, social influence can also be contradictory and confusing. The mechanisms of social learning in groups have not been fully characterized, partly due to the lack of adequate mathematical description. Using the known reinforcing property of social influence we introduce a reinforcement learning model which can account for social effects at the group and individual level. We use it to reveal pairwise influence relations and the overall learning strategies in a reversal learning task applied to a cohort of mice housed in an Intellicage. Animals make an efficient use of social influence when the goal of the group agrees with their own, but switch to an individual learning scheme when these goals are misaligned. They also exhibit varied decision rules depending on their own motivational state, and a selectivity in assimilating social information depending on the state of their conspecifics.

## 1 Introduction

Most animal species share the ability to adapt to their environment through learning. But compared to learning from own experience, i.e. by trial and error, it is better yet to avoid errors by relying on the knowledge of others. The ability to perceive peer influence is not unique to our kind: many animals form groups whose members, through observation and imitation, acquire useful behaviors and make better decisions. But in rich social environments, not everyone has the same goals, and each individual must decide how to cope with noisy and potentially contradicting social information.

Social learning can broadly be classified as observational (vicarious) or imitative, depending on whether outcomes of peers are observable or not [1]. Both phenomena have been studied from the standpoint of many disciplines, but research methods generally fall in two categories. First, we can passively observe spontaneous behavior within social groups and quantify the relevant aspects of it in statistical terms. This descriptive approach is general and applicable to any species, both in wildlife and laboratory, and various learning strategies have been proposed explaining how social influence from multiple conspecifics combines with own experience [2, 3, 4]. However, the complexity of group behaviors and the multitude of confounding factors which can neither be controlled nor accounted for in the natural environment, make it difficult to generalize the facts established in this way. The second, more rigorous approach draws inspiration from the tradition of laboratory experiments started by the studies of Pavlov on classical conditioning and Konorski on instrumental conditioning [5], which established the laws of associative learning by reinforcement, common to human and higher animals. These laws were later formalized as mathematical models [6], leading to the computational reinforcement learning theory [7], the basis of agent-based artificial intelligence algorithms. The advantages of model-based approach over descriptive statistics are prediction and inference: it is useful to be able to simulate the behavior and to reason about the underlying decision making process [8]. Moreover, agent models are an important vehicle in neuroscience because they mimic certain aspects of dopaminergic midbrain activity underlying reinforcement [9, 10, 1, 7]. Dopamine has also been linked with behavioral phenomena such as preference for novelty [11, 12] or perseveration [13], and the reinforcing character of social influence has long been realized [14, 15].

However, the observational nature of learning from examples of others fits poorly in reinforcement learning theory’s underlying formulation of Markov decision process, in which learning is framed as a sequence of trials involving the agent’s action. A widely used experimental paradigm which we adopt in this work is the reversal learning test. The task is to learn the correct choice from among a number of options with the purpose of maximizing reward gained over a sequence of trials while the location of greatest pay-off relocates from time to time (‘reversal’). The reward is immediate and the task has minimal cognitive requirements, relying on the essential ability underlying conditioning, namely, the ability to form expectations about outcomes of stimuli (classical) or actions (instrumental). In consequence, it is applicable to human [16, 10] and other species, including rodents [17, 18]. Reversal learning applied to a single individual reduces to an instance of the multiarmed bandit problem [7], but modelling it in a group poses a greater challenge. Previous experimental studies placed important restrictions on social interaction to make it analytically tractable. Typically, the test is executed only with two individuals (participants) having predefined roles of demonstrator and observer, and with the trials of both parties scheduled in lockstep. These constraints are easiest to impose in human (e.g. [1]) where effects such as reputation can additionally be taken into consideration [19, 20, 21]. Nonhuman primates can be trained to perform the task, both with human and a conspecific as demonstrator [22, 23, 24, 25], whereas in rodents, rules can be enforced by technical constraints [26, 27]. In more ethologically relevant studies involving spontaneous activity of numerous individuals, the model-based approach has remained inaccessible. Thus, our understanding of learning in a general social context is limited.

In this work, we attempt to reconcile both approaches to studying social learning. Our computational model builds on earlier works combining the effect of reward with other reinforcing effects in an action-value choice decision rule [11, 28, 29, 30, 31], including works on social influence in constrained interaction regime [1, 29, 32, 20]. Rather than cramping interaction, we generalize the formulation of social influence to make it applicable to spontaneous activity. We use our model to inquire into the learning strategies in a cohort of mice housed together in an Intellicage [33, 34]. The Intellicage has previously been used for assessing the phenotype in terms of learning [35] and social behavior [36] and to study the interplay between these terms [37], but usually through the descriptive approach. Of specific interest in our study is the role of goal sharing. In our experiment, reward is not always collocated with the rest of the group. Using our model-based approach, we show how the decision making process changes when social influence, so far helpful, suddenly becomes confusing. We also characterize how influence on the individual level shapes the structure of the group. Our results indicate that mice maintain separate value-based decision rules depending on their motivational state and selectively integrate the observed choices of others. Our companion paper [38] gives a broader treatment of the learning behavior, including its temporal dynamics and a family of learning models, in a setting with mean (i.e., collective) social influence signal.

## 2 Results

### 2.1 Social experiment

Our experiment was designed to expose social influence in a deterministic reversal learning task (see Methods). A cohort of 14 mice was housed together in an Intellicage (Fig. 1A). Animals could drink in the four corner chambers of the cage, each containing two bottles, one with plain water and another with saccharine solution (reward). Access to the bottles was controlled by computer-operated sliding doors mounted on nose ports on both sides of the chambers. At all times, only one bottle was accessible in such a way that reward was always offered in one corner and water in the remaining three. To maximize their gain of the sweet reward, animals had to continuously learn the correct location which changed every 2 days.

**Figure 1:**
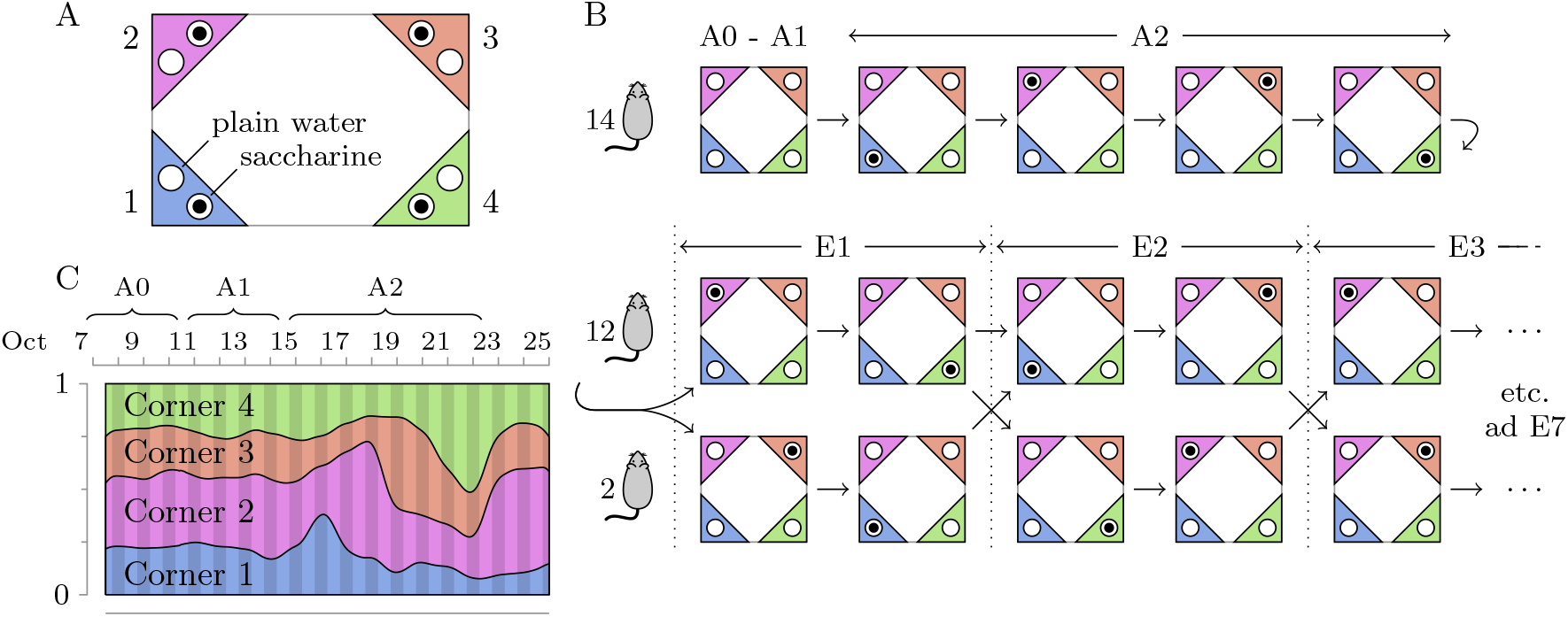
A, Schematic depiction of the Intellicage and the placement of bottles in corner chambers. B, Reward schedule over the course of the experiment (simplified cage pictograms showing only accessible corner sides). Following initial adaptation (top row), the main experimental part started (phases E1 to E7, truncated at E3). In each phase, two animals were assigned minority roles with reward located in different corners than the remaining 12 animals (majority). The reward sequence (2-4-1-3 for majority, 3-1-4-2 for minority) repeats after every 2 phases. C, Proportion of corner visits over the course of adaptation phases, averaged over all mice. A spontaneous overall preference for Corner 2 develops in the group early in the experiment. Conditioned preferences, coincident with reward, superimpose on the generally constant spontaneous proportions.

Upon completion of 16-day adaptation (phases A0 to A2, top row in Fig. 1B) in which reward was first introduced, the main part of the experiment started (phases E1 to E7, truncated at E3 in the middle and bottom rows in Fig. 1B). Animals were divided into two groups, of which one counted 2 animals (‘minority’), while the other (‘majority’) consisted of the remaining 12. Each group was rewarded in a different corner. This was achieved by exercising per-animal selective access control to bottles, conditional on the visiting animal’s current group assignment. Assignments changed from phase to phase in a 4-day cadence, in such a way that each animal was part of the minority group in exactly one phase in the course of the experiment. Thus, both the majority and minority corners served saccharine solution to animals from the corresponding group, but water to the opposite group. As a result, the learning objectives of animals from the two groups differed.

As expected, saccharine has a positive reinforcing value and drives corner preference variation, as shown in Fig. 1C, in which the proportion of corner visits was estimated using a cosine tapered 24-hour window moving in time (an initial time interval is shown covering adaptation phases A0-A1 without reward and A2 with reward common to all animals).

Compared to reward, the effect of social influence is more subtle. As a preliminary check, we leverage the design of experiment to see if we can detect a propensity to visit corners rewarded for, and thus preferred by, the opposite group in phases E1 to E7. To verify that the majority influences the choices of animals during their minority assignment (such influence can reasonably be deemed stronger than in the reverse direction due to group imbalance), we compare the proportion of visits of minority animals to corners rewarded for the majority with these same corners during the phases in which they were not rewarded for either group (neutral). We estimate proportions in the second halves of phases E2 to E7, as the initial halves of these phases located the group’s reward in the same location as of the majority at the end of the previous phase (i.e., minority animals did not initially experience reversal and knew the correct corner); the first half of E1 was used as this phase as the above circumstance did not apply. Paired t-test reveals a significant positive effect, resulting in average visit proportion increase from 0.1822 (neutral) to 0.2362 (*t*_13_ = 3.016, *p* = 0.009921, two sided). The influence which the minority exerts on majority is less pronounced but nonetheless significant and positive: the average proportion shifts from 0.1793 (neutral) to 0.2169 (*t*_55_ = 4.679, p = 0.00001917, two sided). While the effect is weaker, the power of the test is increased since, during majority assignment, proportions can be estimated and compared for each corner rather than just one. Observe that, absent social influence, we should expect negative dependence, stronger in the minority condition, as a result of greater corner congestion.

### 2.2 Corner choices are driven by multiple factors

We now focus on model-based description of social influence as a conditioning stimulus acting in parallel to reward. Our approach builds on the established probabilistic model of learning in which the probability of choice *c* ∈ 𝒞, in our problem distributed over the four cage corners 𝒞 = {1, 2, 3, 4}, is conditioned on the history *H*_*n*_ of events which had happened until trial *n*

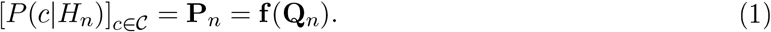

The second equality links the probability distribution in trial (visit) *n*, encoded as a vector **P**_*n*_, with the vector of corner values **Q**_*n*_, using a multivariate logistic mapping. The value vector **Q**_*n*_ aggregates the history in a markovian way. In our model, this quantity is constructed as a linear combination of three kinds of valuation components, each representing a different aspect of the history: the reward, the others’ observed choice and own choice.

As a prerequisite for further considerations, we introduce a convention to uniquely name quantities relevant to individual animals. The *focal animal* is the animal which we model, denoted *i*, while *j* shall indicate any other *influencing animal* whose behavior impinges on the focal. Let *R*_*in*_ ∈ {0,1} and *C*_*in*_ ∈ 𝒞 denote, respectively, the reward and choice of the focal, while *Cj*_*n*_ shall stand for the corner choice of another animal *j*, where in our case *i, j* ∈ 𝒜 = {1, …, 14}. The focal-specific rewards and choices are considered to form contiguous sequences (*R*_*i*>*n*_) and (*C*_*i*>*n*_) for n = 1,…, *N*_*i*_, even if in reality they are interspersed by visits of other animals.

First, we model the conditioning effect of saccharine using Rescorla-Wagner update rule [6]

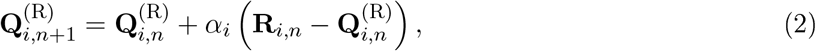

expressing the concept of value (reward expectation) changing only as a result of directly experienced outcome. The change is proportional to the reward prediction error (RPE), or the difference between expected and experienced reward value. We employ the fictive update scheme [39, 16], adapted to multiple choice situation. On each trial, values of all options are updated, not just the one selected, reflecting the idea that both positive and negative outcomes can become informative once animals develop an expectation of reward. We encode the elements of the reward vector **R**,_*i,n*_ as *R*_*i*, *n,a*_ = *R*_*i*, *n*_ for *a* = *C*_*i, n*_ and 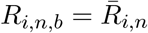 for each *b* ≠ *C*_*i, n*_, where 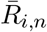 represents a fictive reward value, as opposed to the factually experienced *R*_*i, n*_. To balance correct and incorrect choices, we set 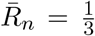, giving equal credit to the three remaining options when reward is missed. This update scheme is a simplified alternative to baseline adaptation [7], avoiding unnecessary complication of valuation dynamics.

Next, we model social influence of animal *j* (*i* ≠ *j*) using a valuation rule analogous to (2)

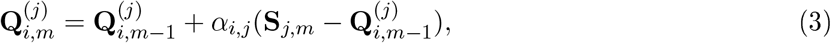

but based on social influence signal **S**_*j*,*m*_, a one-hot encoding of *j*’s choice. It bears resemblance to reward in the fictive setting, where each observed choice of others is perceived as a positive outcome. Such a definition is adequate in imitational social learning and corresponds to the fact that Intellicage corner chambers can harbor one mouse at a time, in principle making no information about saccharine available to individuals outside. The method is referred to as observational action prediction error (APE) [1] as the social valuation 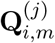 tracks the current proportion of visits of animal *j* in each of the corners. A new visit index *m* reflects the fact that, in spontaneous behavior, the social valuation sequence is temporally unaligned to the sequence of focal’s visits and thus cannot be combined with reward-based valuation. Previous studies in human [1, 29, 20] enforced alternation of demonstrator and observer choices to achieve aligned value signals, which cannot apply in our case. Instead, we introduce the time alignment operator, see Equation (9) in Methods, which can be interpreted as sampling from a zero-order hold interpolated valuation, as visualized in the plots on the right in Fig. 2. The underlying principle is that influence-derived values do not change in between the times of visits of the animal from which they originate, consistent with the dynamics of valuation in Rescorla-Wagner model. The result of applying this operator to the social valuation 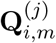 is a time-aligned counterpart 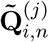 as perceived by the focal at the time instants of decision making.

**Figure 2:**
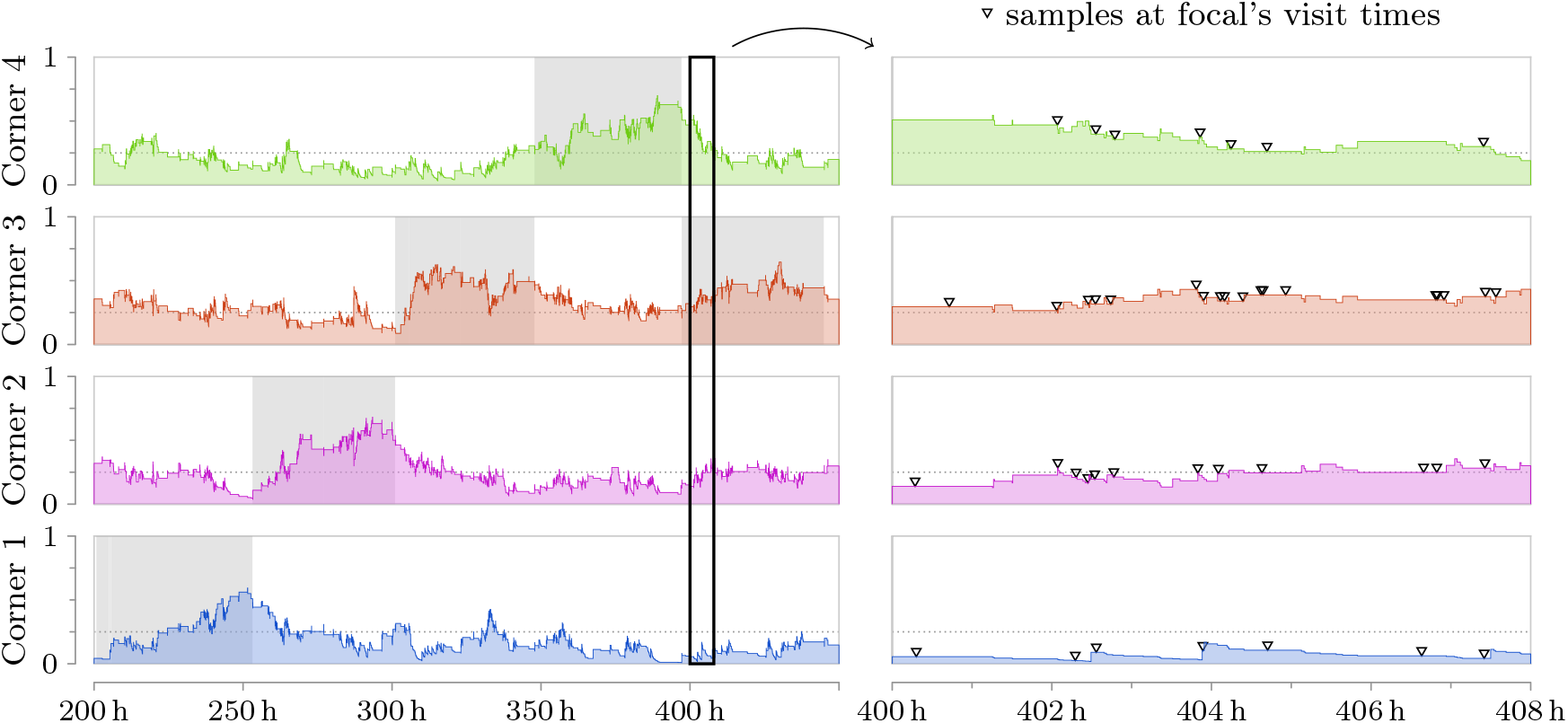
Social effect of Animal 1 (**Q**^(1)^) plotted against real time, with *α =* 0.0736 fitted to Animal 2 as focal. Color-coded plots show the interpolated piecewise constant valuation signal, Eq. (3), in each of the four corners. The time range of the plots on the left covers the total of adaptation phase A2 and the beginning of E1 in which both animals are in their minority assignment. Shading marks periods in which respective corners were rewarding. The plots on the right illustrate the concept of time alignment within the time range marked by the bold rectangle. Following saccharine relocation, the observational value of Corner 4 declines while values of Corner 3 (rewarded for Animal 1 and 2) and Corner 2 (rewarded for the majority) increase.

Finally, we include a term based on own action prediction error, which we refer to as *automodulation*. Its valuation formula is identical to (3) except it is calculated from the history of own choice, by posing focal’s index *i* for *j*:

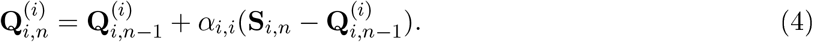

A value-free reinforcing mechanism of this type is known in the literature as the habitual system [40, 41]. It was proposed to explain the emergence of habits, defined as a tendency to repeat actions often selected in the long run regardless of whether they are rewarded or not. This tendency had been reported in many earlier behavioral studies, including in mice [42], and attributed to perseveration [43, 30, 44] or described as “stickiness” [18] or choice autocorrelation [29]. We chose ‘automodulation’ as a more general term to reflect the fact that the reinforcement based on own action prediction error can be both positive and negative, as described later.

In its full generality, our model accounts for social influence from each individual in the group, which leads to a total of 15 effects per focal animal in our cohort, including automodulation and reward (by ‘effect’ we mean valuation originating from a separate RPE or APE signal). Each effect is associated with a pair of parameters: the learning rate and strength (*α*_*i*_, β_*i*_ for reward and *α*_*i,j*_, *β*_*i,j*_ for social influence and automodulation). The effect strengths are the scaling coefficients in the linear combination formula for the net option values (Equation (10) in Methods), from which choice probability distribution is obtained. Additive construction of action values from component effects has been employed in prior studies [43, 30], including in the context of social learning in a constrained setting [1, 29]. We name models based on this principle *plurivalent*, in contrast to models where valuation is based on a single effect, which we refer to as *univalent*. The univalent reward model has been widely used in studies on reinforcement and constitutes a useful reference. It is instructive to study the dynamics of this and other effects, both in isolation and in combination.

Fig. 3 shows log-likelihoods or their two dimensional sections through the parameter space of the effect of interest for five selected animals. Column A shows univalent models of reward. Except for Animal 13, all animals in the cohort display a stereotypical likelihood shape attaining a single peak with *α* in the range from 0.05 to 0.1, indicating a relatively slow learning of reward based value, and *β* around 2. This contrasts with the theoretically optimal win-stay, lose-shift policy corresponding to *α* = 1 and *β* → ∞ to. However, this theoretical result requires the knowledge of experiment design, which mice obviously do not have. Their behavior, as interpreted based only on the history of reward, shows a balance between exploration and exploitation of the already acquired knowledge about the environment. In Animal 13, likelihood peaks at strongly negative *β* and a very low *α*. This, in our view, reflects the effect of competition for rewarded corners rather than avoidance of saccharine. We reject this maximum and find a negligible secondary log-likelihood peak of 0.072 at *α* = 0.8, *β* = 0.05, revealing inability to learn from own experience.

**Figure 3:**
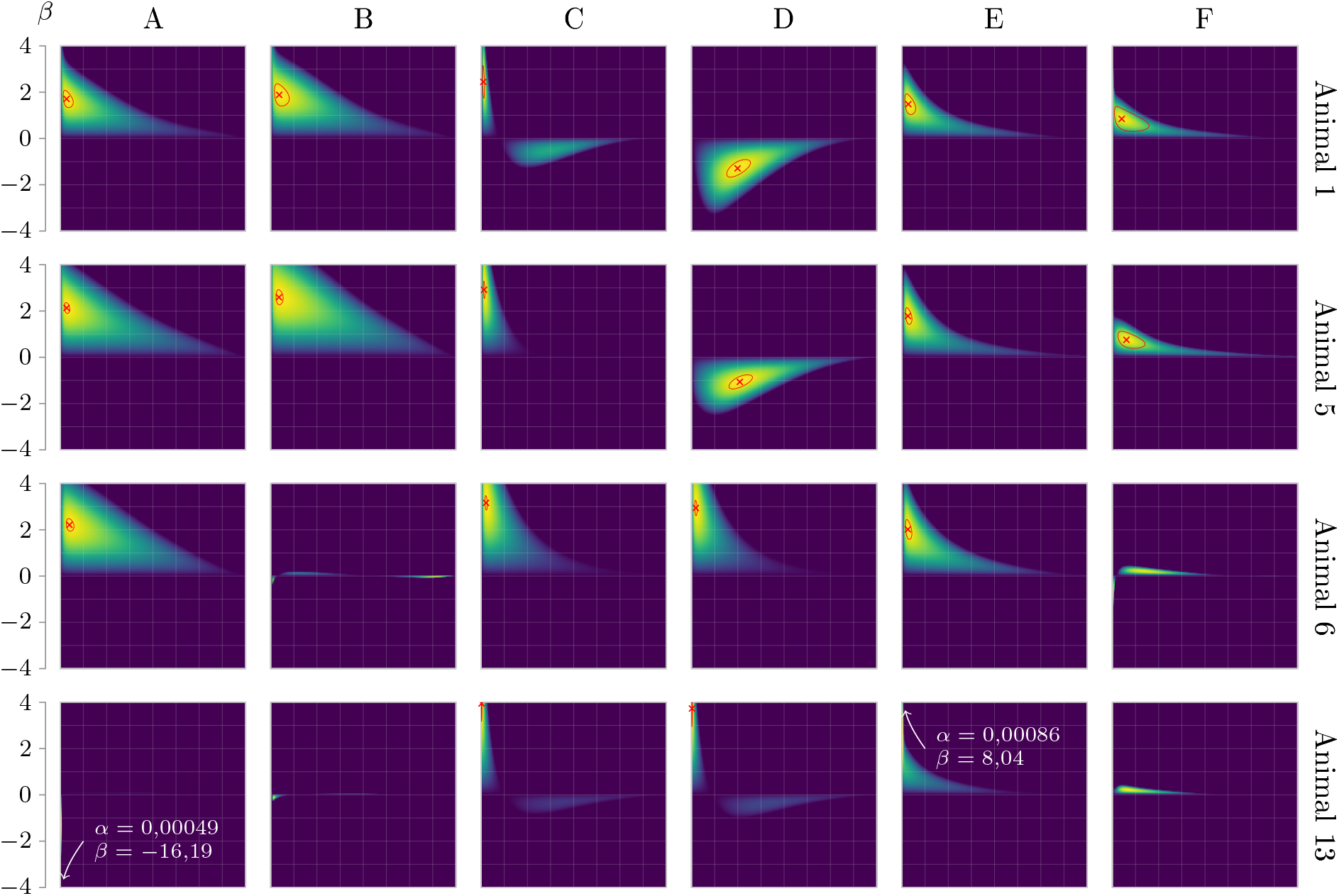
Comparison of log-likelihood functions in the parameter space of the effects of interest for five selected focal animals (arranged in rows). A, univalent reward model; B, section through reward parameters in a bivalent reward + automodulation model; C, univalent automodulative model; D, section through automodulation parameters in the same model as B; E, univalent model of influence of Animal 11; F, section through the parameters of the influence of Animal 11 in a plurivalent model with BIC-selected effects. All plots share *α* and *β* scales and the surface color scale ranges from reference (darkest) to model maximum (brightest). Reference level is the null model for univalent models (A, C and E) or the level of the stated model with effect of interest removed. MLE points and the surrounding 95% confidence regions are marked in red for effects selected by the BIC criterion.

Peak log likelihood levels relative to null model (RLL), corresponding to the models from Fig. 3 A, are presented as red bars in Fig. 4 A. As a complementary measure independent of the number *N*_*i*_ of visits of the focal, we use per-trial prediction accuracy which estimates the probability of correct prediction of choice in a single trial given the history (an optimistic estimate because based on reclassification of data used in fitting, see Methods). Accuracy values of univalent reward models fitted to our data are modest (supplementary Fig. 10), staying generally below 0.3 (with chance level of 0.25). However, when fitted to the subset of visits in which licks were detected, reward models yield markedly greater likelihoods (blue bars in Fig. 4 A) and accuracy in excess of 0.4 in 7 of 14 animals. In visits without drinking (‘idle’, which make ca 50% of cases), mice have no gustatory sensation of outcome, thus in principle a model of reward-based conditioning should be equivalent to random choice. We still observe fits with appreciable RLL in some animals (green bars) and implausible MLE parameters, *α*_*i*_ ≈ 0 and *β*_*i*_ < 0, which we again attribute to corner congestion (models fitted under *β*_*i*_ ≥ 0 constraint are indeed characterized by RLL scores not significantly greater than zero). These results show, first, that the decision to drink is primary with respect to corner choice, and second, that animals carefully select the corner for drinking, certainly based on reward history. The poor prediction accuracy of the model fitted to complete data apparently results from the inclusion of idle visits in the history, which seem to be irrelevant for RPE valuation and thus only contribute noise. Nevertheless, in an attempt to explain the decision making process as a whole, we consider drinking and non-drinking visits together in the rest of the text, unless otherwise stated.

**Figure 4:**
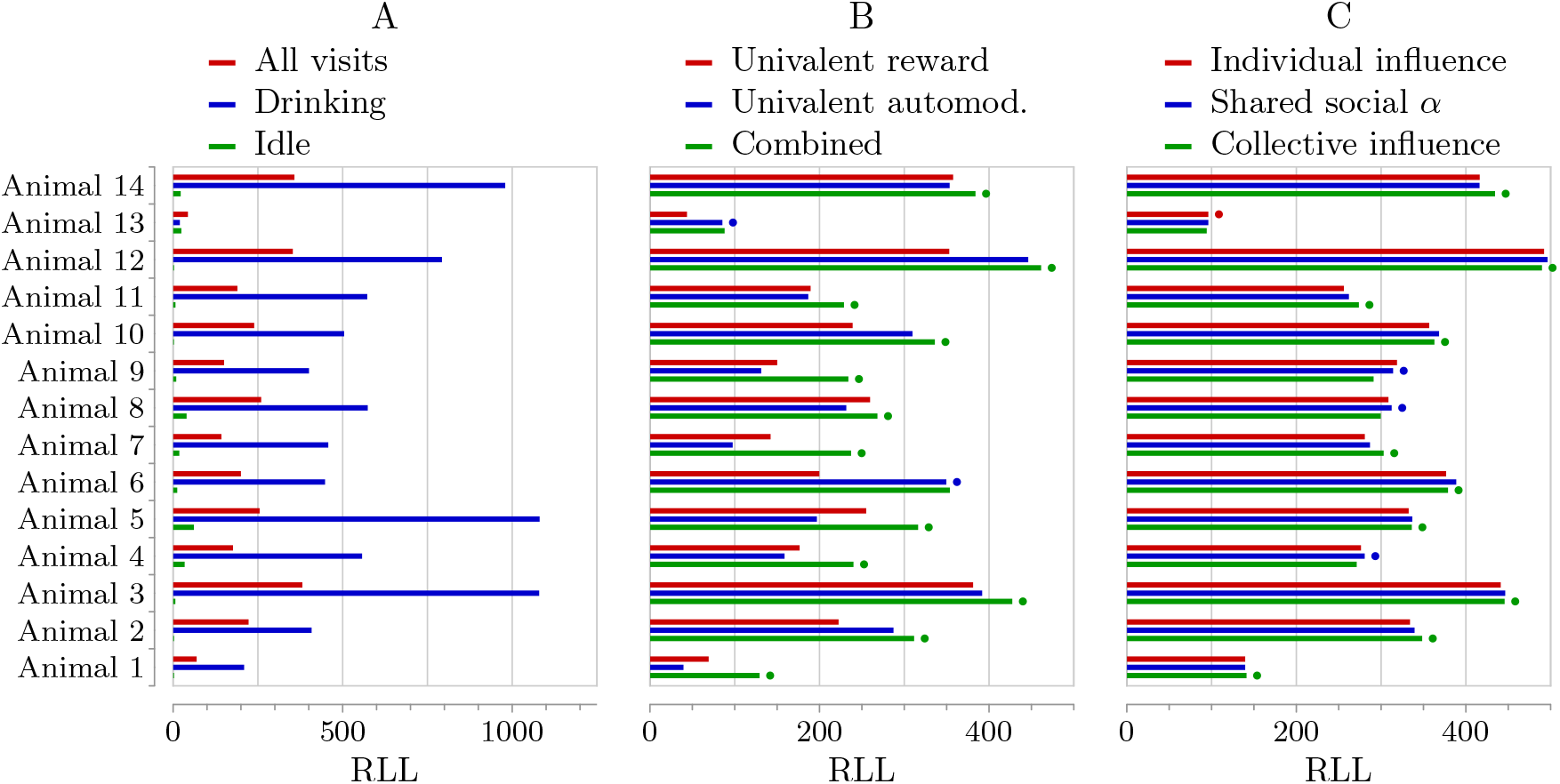
Likelihood comparison of models. A, reward models fitted to data subsets defined by lick condition in phases E1–E7; B, isolated reward and automodulative effects and their combination (all visits in phases E1–E7); C, plurivalent models with selected effects and varied constraints on social influence representation (all visits in phases E1–E7). Dots mark the model selected by BIC.

Plots in Fig. 3 C illustrate the dynamics of the isolated automodulative effect, which is qualitatively different than the reward. The positive RLL wedge, which for reward narrows gradually as *α* → 2, shrinks abruptly and above an individual threshold, the automodulative effect either remains negligible or, in some animals, we observe a secondary likelihood maximum in the negative *β* halfplane. The positive automodulation (*β*_*i,i*_ > 0) corresponds to the habitual system, but negative automodulation (*β*_*i,i*_ < 0) is qualitatively different, and has not been reported before. The fact that it comes with considerably greater learning rate, corresponding to a short time scale, allows it to be interpreted as a preference for (relative) novelty.

Reward and automodulation change their valuation dynamics when combined in a bivalent model, depending on which of them is a better univalent predictor (cf. Fig. 4 B). This is depicted in columns B and D in Fig. 3. When the effect of reward dominates automodulation (Animal 1 and Animal 5), the latter undergoes a qualitative change: the likelihood peak in the positive *β* halfplane disappears and a single peak remains in the negative *β* halfplane. This may happen even if no signs of bimodality are seen in the univalent automodulation plot, as is the case in Animal 5 in Fig. 3 C. On the other hand, when positive automodulation dominates reward, the latter is suppressed, sometimes completely (Animal 6 in Fig. 3 B). This illustrates the risk of false inference posed by underspecified models. In view of BIC comparison (Fig. 4 B), the apparent conditioning effect of reward in Animal 6 turns out spurious, and only automodulation matters. In Animal 13, the effect of reward is negligible and the bimodal dynamics of automodulation is retained. Barring this exceptional case, bivalent (reward + automodulation) models display generally unimodal dynamics where automodulation is either positive (habitual) or negative (a tendency to chose options long unexplored). They also attain RLL scores often greater than either of its constituent effects taken separately (Fig. 4 B).

The plots in Fig. 3 E show examples of log likelihoods of univalent models of social influence of Animal 11. The isolated effect of social influence displays dynamics similar to reward. Indeed, the choices of others can reveal the information about reward location when the latter is not represented in the model, as shown in the valuation plots on the left in Fig. 2. But social influence is a useful predictor also when the conditioning effect of reward is properly accounted for in the model (Fig. 3 F). Strong influence of Animal 11 is visible in Animal 1 and Animal 5, for which this effect is selected as a relevant predictor in a selection procedure described later, while in remaining focals it is eclipsed by other competing effects.

### 2.3 Scaling the model

We now consider what all effects taken together can tell us about the mice and how the model defined in Methods can be tamed. With *K* = 14 interacting mice, the full model of one focal has 15 effects and 30 parameters. Likelihood maximization in a parameter space of this dimension is nontrivial as the optimization problem is nonconvex. In this section, we consider ways to simplify the model and facilitate domain-wide likelihood maximization.

To start, we introduce the constellation plot, a convenient way of visualizing plurivalent models as exemplified in Fig. 5. The *constellation* is a configuration of point estimates of component effects, called *effect loci*, in a common (*α, β*) plane. The coordinates of each locus are the learning rate and strength of the corresponding effect. Shaded areas surrounding the loci depict marginal 0.95 confidence regions of parameter estimates. For ease of inspection, we consistently mark social effects as blue dots, automodulation as red crossed circles and reward as black crosses.

**Figure 5:**
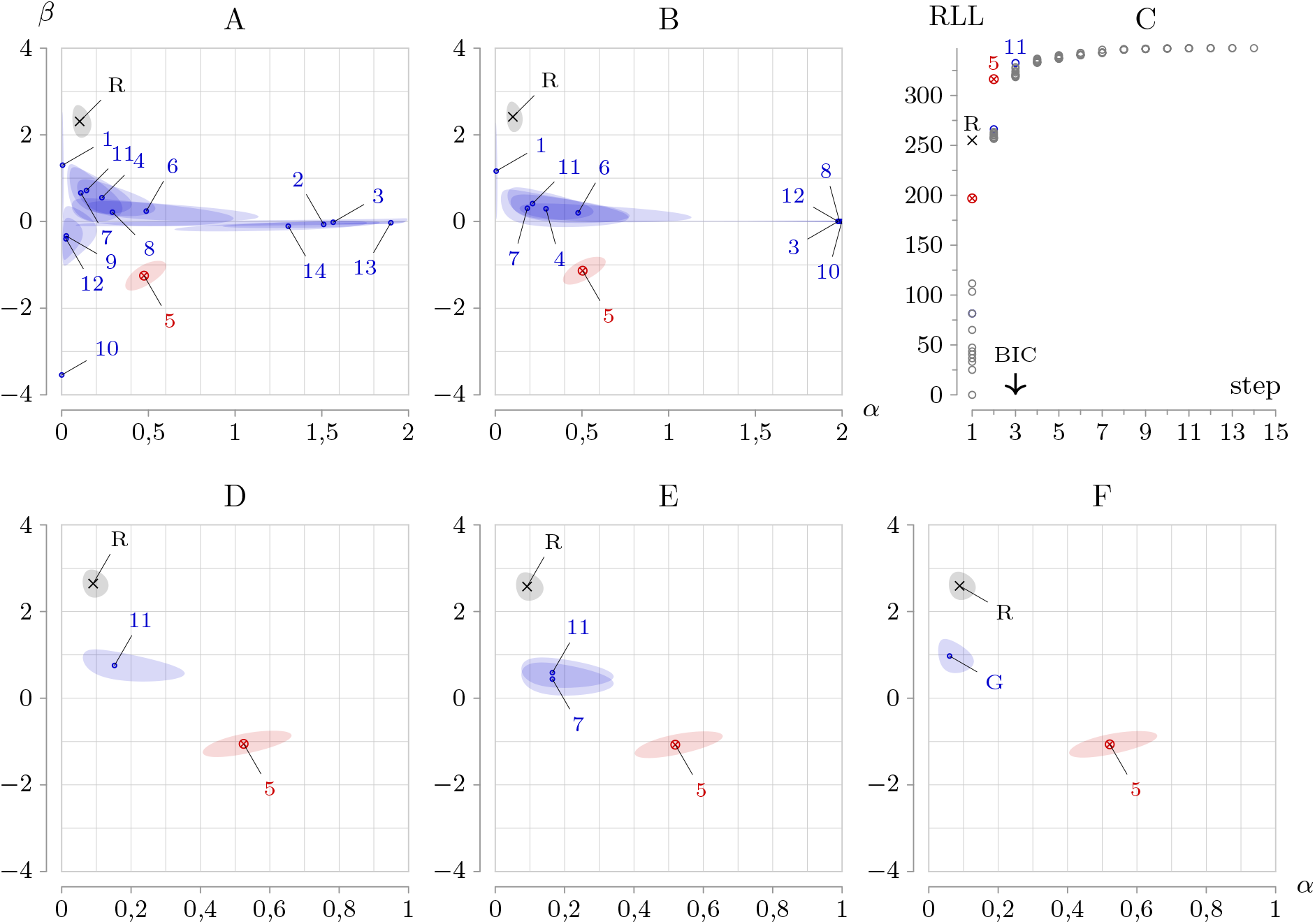
Constellation plots of models of Animal 5 and model selection. A, full order model encompassing all effects with no fitting constraints; B, full order model encompassing all effects with positivity constraint on social weights; C, illustration of the forward stepwise selection, with selected effects highlighted and remaining grayed out; D, reduced model with three effects selected by BIC as shown in C; E, reduced model with shared social learning rate (one additional social effect is selected); F, reduced model with collective social effect. Numeric labels refer to the enumeration of individual mice, R − reward, G − group (collective effect).

The constellation in Fig. 5 A, besides being overly complex, reveals problems with the model fit. First, some loci cluster along the *α* = 0 axis, sometimes with negative *β*. These long-term social effects reflect similar corner preferences rather than influence. Following the rationale given in Methods, we impose the constraint *β*_*i,j*_ *≥* 0, except for automodulation (Fig. 5 B). Second, we restrict the model to only include the subset of important effects through a stepwise selection procedure whereby effects are fitted incrementally and their representative subset is selected using the Bayesian Information Criterion (BIC, see Fig. 5 C). Through this, we remove a considerable fraction of weak effects with *β* ≈ 0 (mainly social, but reward and automodulation also turn out insignificant in some animals), which are prone to overfitting to an arbitrary *α* value. Our approach is heuristic, but based on objective criteria. While giving no guarantee of success, it yields reasonable models in our cohort when fitting to the complete main experimental phase, without resorting to regularization which we reserve only for difficult cases such as fitting to short observation periods. For details, see Methods.

When considering social effects, one may argue that the learning rates *α*_*i,j*_ relate to memory processes rather than having a social character. According to this account, only *β*_*i,j*_ parameters are allowed to vary by influencer but not the learning rate: *α*_*i,j*_ = *α*_*i,k*_ for *j, k* ≠ *i*. As a result, social effects starting with the second cost only a single new *β* parameter and model selection based on BIC favors richer descriptions. This variant (called ‘shared social *α*’) is illustrated in Fig. 5 E, where one additional effect (Animal 7) is included in the constellation.

As a further simplification, we can assume social influence not distinguishing individuality but having an aggregated (mean field) nature, similar to [38], by considering visits of others as one signal originating from a single source, the group. The model variant fitted under this assumption is shown in Fig. 5 F. Compared to individual influence, the learning rate of collective influence is in general smaller to compensate the increase in frequency of visits of the group (approximately 13 times greater than individual).

Likelihood and BIC comparison (Fig 4 C) does not give a decisive indication of which of the three variants of social influence best explains behavior. As a rule, influence with shared *α* is more economical in terms of model degrees of freedom. Consequently, BIC tends to select a richer set of social effects, leading to greater likelihood (blue bars) compared with the variant in which all parameters are free (red bars). The collective influence of the group, on the other hand, is a signal with more observations and thus potentially less noisy. This makes it a strong predictor in most, but not all, focal animals (green bars). We give an interpretation for this fact later. The generally small log-likelihood differences between the three variants allow their interchangeable use, tailored to specific needs. We proceed by giving two examples.

### 2.4 Pairwise influence reveals the group’s structure

The graph shown in Fig. 6 A shows the map of social influences across the group, determined using models with shared social learning rate and selected subset of effects, fitted to all visits in phases E1 to E7 for each animal. Influence arrows are directed from influencer *j* to focal *i* and their widths reflect *β*_*i,j*_ (these parameters are all positive owing to the constraint in the fitting procedure). Mutual influence is detected in some pairs of animals as indicated by broken arrows pointing in both directions. Most arrows are, however, unidirectional, which is consistent with our definition of influence as a causal and directed relation. Judging from the number of detected social effects, mice differ in their social rank. Animal 7 and Animal 11 seem to be the most influential with 6 and 5 oubound arrows, respectively. On the other extreme we find Animal 8 with no detected effect on peers. Because social learning rates are in general different between focals, only the strengths of inbound influence can be compared, but not of outbound. Moreover, subset selection leads to slightly overestimated social effects (since they must compensate for the rejected effects). In consequence, strengths of social effects do not constitute a good measure of social influence.

**Figure 6:**
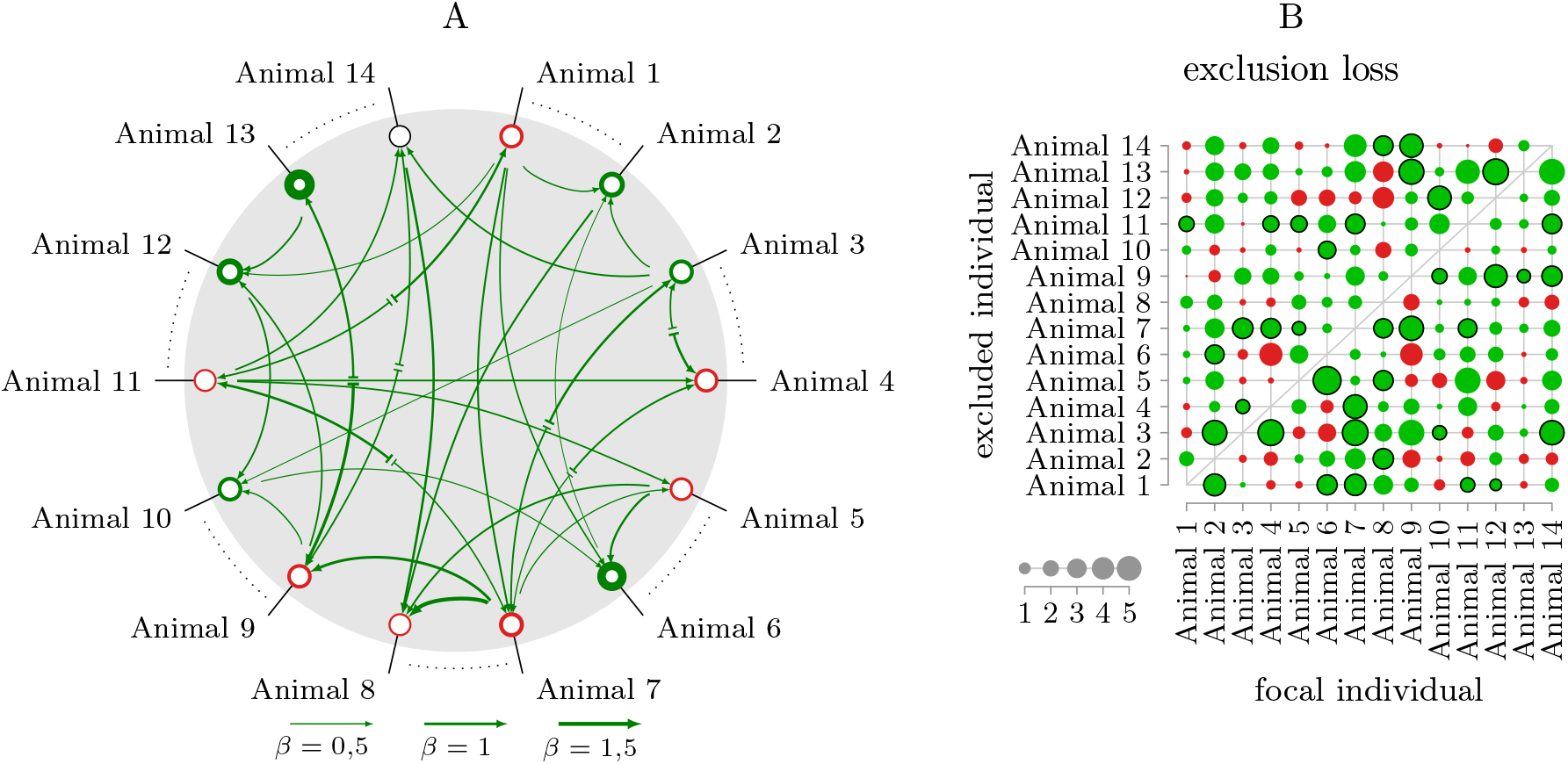
A, Social influence network obtained by combining models fitted to individual animals as focal, with shared social a parameters. Line thickness of incoming arrows represents the weight *β*_*i,j*_ of influence of *j* impinging on focal *i*. Green and red rims surrounding animal nodes depict, respectively, positive or negative automodulation weight, using the same line thickness scale. B, Exclusion loss in the collective influence model (green – positive, red – negative); dots with black outline indicate effects included in the graph in A.

A reasonable approach to evaluate the magnitude of a social effect is based on a likelihood comparison between two models, one including the animal of interest and another in which it is excluded. To avoid confounding model degrees of freedom in the comparison, which would occur if influences were represented as individual effects, we base the comparison on the collective influence variant of the group in two models, one reference and the other model fitted to data with visits of the animal of interest removed (see Methods). The resulting measure, which we call *exclusion loss*, reveals how much the predictive power of the collective effect relies on the animal of interest.

Fig. 6 B shows exclusion loss for each pair of animals. It is not an absolute measure as in some cases it takes on negative values (red). In these cases, the exclusion of the incluencing animal improves prediction, rather than worsening it, indicating a weak or no influence. This supports our hypothesis that mice differentiate the influence from conspecifics. The exclusion loss is also not symmetric, consistent with the expectation that social influence is a directed relation. To compare animals in terms of their social rank, we can sum the exclusion loss over focals (in each case, it is a different set). The aggregated values confirm the dominant roles of Animal 7 and Animal 11, but also Animal 3 and Animal 13, and the lowest rank of Animal 8.

It is noteworthy that the subset of social effects depicted in the graph in Fig. 6 A corresponds to robust effects as measured by exclusion loss. In Fig. 6 B, the selected subset of these social effects is additionally marked by a black outline. In general, model selection does not guarantee chosing effects in order of their magnitude, since effects already included in the model tend to reduce the impact of most closely linearly related candidates in the current step. Nevertheless, we see that in each focal, the subset of social effects is chosen from among the strongest influences.

Finally, we refer back to Fig 4 C and propose the explanation for the fact that some animals are better explained by a subset of individual social effects, while others by collective influence. We attribute this fact to the variation of magnitudes measured by exclusion loss within the focal of interest, as shown in Fig. 6 B. Animals with more uniformly distributed exclusion loss seem to attain greater likelihood in the collective influence model. Conversely, more socially selective animals are better modelled by models of individual social effects.

### 2.5 Motivational state affects behavioral patterns

We now attempt to interpret the behavior of mice from the constellation plots shown in Fig. 7, in which collective influence is used for simplicity. The models in row A were fitted to all visits in phases E1 to E7. Automodulation loci are present in all models, but the associated weight parameter *β*_*i,j*_ is positive in half of the cohort (7 animals), and negative in the remaining half. The learning rate *α* is generally small in positive cases, indicating long-term habitual dynamics, while in negative cases it is much greater. The collective influence of the group emerges in all models as a positive effect, showing a tendency to choose popular corners. The reinforcing effect of reward is not ubiquitous. It is absent in Animal 13 and Animal 6, in line with earlier results in the bivalent reward + automodulation model.

**Figure 7:**
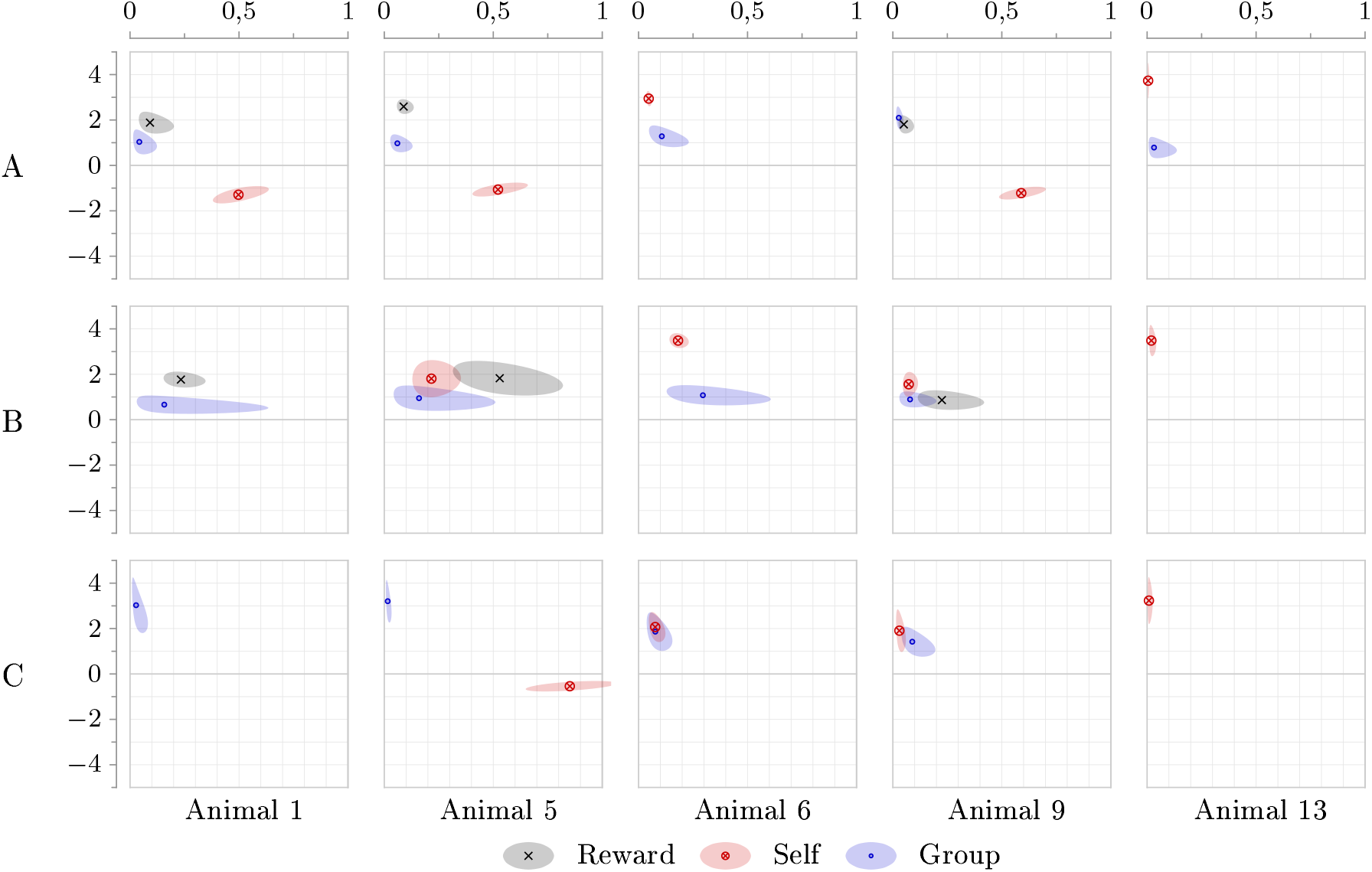
Constellation plots representing models of chosen mice fitted in phases E1–E7 (see supplementary Figs. 11–13 for the complete cohort). A, Unconditional case (all visit data); B, goal-oriented case (only drinking visits); C, idle case (only non-drinking visits).

Our earlier analysis suggests the existence of two motivational states that differentiate the strength of reward as predictor of choice and which we term “goal-oriented” and “idle”, corresponding, respectively, to the drinking and non-drinking condition. To see how social learning depends on these states, we now scrutinize plurivalent models fitted separately in either condition. Note that state-relevant social influence can be modeled in two ways: based either on the full set of observed choices or a subset similarly defined by others’ drinking condition, the latter implying the ability to determine the motivational state of others. Likelihood comparison of models with social signals specified either way gives a decisive support for the latter possibility. Dependent samples t-test confirms systematic likelihood increase both in focal’s idle condition with social effect based on idle visits (mean RLL difference 14.9, *t*_13_ = 3.54, *p =* 0.0036, two sided) and in focal’s goal-oriented condition with only goal-oriented visits observed (mean RLL difference 5.64, *t*_13_ = 4.22, *p* = 0.001, two sided), as compared to unconditional observation models. We conclude that mice are in general able to recognize the motivational state of their conspecifics and selectively imitate their behavior depending on how it relates to own motivational state. One exception exists: Animal 2 in which unconditional observation model wins by 14.45 in idle and 6.86 in goal condition.

The constellation plots in row B in Fig. 7 depict regularized models fitted in goal-oriented state, whereas row C shows the idle state. In all these models, we use the subset of observed visits matching the drinking condition of the focal as the social signal. State-specific models (rows B and C) do not simply combine to unconditional all-visit models in row A. Automodulation has positive (habitual) dynamics in all animals in the goal oriented state (supplementary Fig. 12) and is likewise positive in idle state in all except five animals (supplementary Fig. 13). Two such exceptional cases, Animal 1 (no automodulative effect in idle condition) and Animal 5 (negative automodulation in idle condition), are depicted in Fig. 7. In the remaining cases, automodulation is positive in both conditions even if all-visit fits show negative dynamics (e.g. Animal 9).

To explain why, by combining data subsets defined by drinking condition, each showing a habitual effect, we can arrive at models with negative automodulation, we set forth the following hypothesis: mice maintain independent valuation processes in each motivational state. An individual in a given motivational state makes its choice based on the current values corresponding to that state and reevaluates them based on the visit outcome. Animals intermittently switch between states, and thus alternate between two distinct behavioral patterns. This alternation is the explanation for the negative dependence on choice history observed in some animals, when behavioral patterns differ.

If the premises of the hypothesis held and corner preferences did build up independently, then we should observe within-individual differences in the distribution of corner visits between the two conditions. Indeed, proportions of corner visits calculated over phases E1–E7 separately from idle and goal-directed subsets show substantial differences in some mice (supplementary Tab. 1). Moreover, animals in which visit distributions show greatest discrepancies between conditions correspond to negative estimated automodulation effects, whereas those in which differences are less pronounced have positive automodulation effects, which agrees with our conjecture.

It is noteworthy that the social effect emerges as an important factor across both conditions and in all mice, except Animal 13. The hypothesis that valuation processes are independent in each motivational state gives us a basis for interpreting how social effects vary between the two conditions. In the idle state, social influence tends to be a long-term effect. In comparison, loci of social effects in goal-oriented state are often shifted right, indicating short-term dynamics. This difference can be attributed to the fact that social influence is perturbed when animals’ majority-minority assignments change in the course of time. This perturbation can be expected to impinge on goal-oriented social valuation, where reward constitutes a benchmark to which the usefulness of social influence is referred. No such reference exists in idle case and thus it should not be affected by changes in group assignments, allowing it to develop as a long-term effect.

In the goal-oriented state, the effect of reward is again missing in Animal 6 and Animal 13, and is positive in all remaining mice. In idle condition, reward is excluded from the models based on our prior knowledge. This prevents possible fits of reward effect with negative *β* and very small *α*, reflecting corner congestion.

### 2.6 Mice alter their learning strategy when social context fails

Until now, we considered the decision making process as stationary and treated our model parameters as constants. That is, we assumed that the learning strategy is insensitive to majority/minority assignment and that the animals had developed, by the end of the 16-day adaptation period, a stable social structure and become familiar with the rules of the experiment, expecting reward in a changing location. This assumed stationarity need not hold in general. We are now interested in how the minority assignment affects behavior. Positive social influence obviously helps when the goals of the focal and other mice are aligned, but in minority assignment it is confusing. In this case, the focal animal can continue using its current strategy (probably resulting in less saccharine gained) or change it somehow in an attempt to improve its decision making process. To assess this, we apply the model with collective influence to data in goal-oriented condition using interval-focused weighting (see Methods), obtaining model fits tuned to behavior in the period before, during and after minority assignment. As we only consider phases from E1 to E7, we cannot estimate the pre-minority condition in animals 1 and 2, likewise we have no data for post-minority condition in animals 13 and 14, which we remove from the pool of models. We also remove animals 3 and 4 due to a considerable gap in data in phase E2 caused by technical problems with the Intellicage. Fig. 8 presents the constellations corresponding to the three situations in the remaining eight mice. The models show effects selected in the usual procedure, except regularization was additionally employed to suppress artifacts caused by insufficient fitting data, especially in minority assignments where the fitted period covers only one phase.

**Figure 8:**
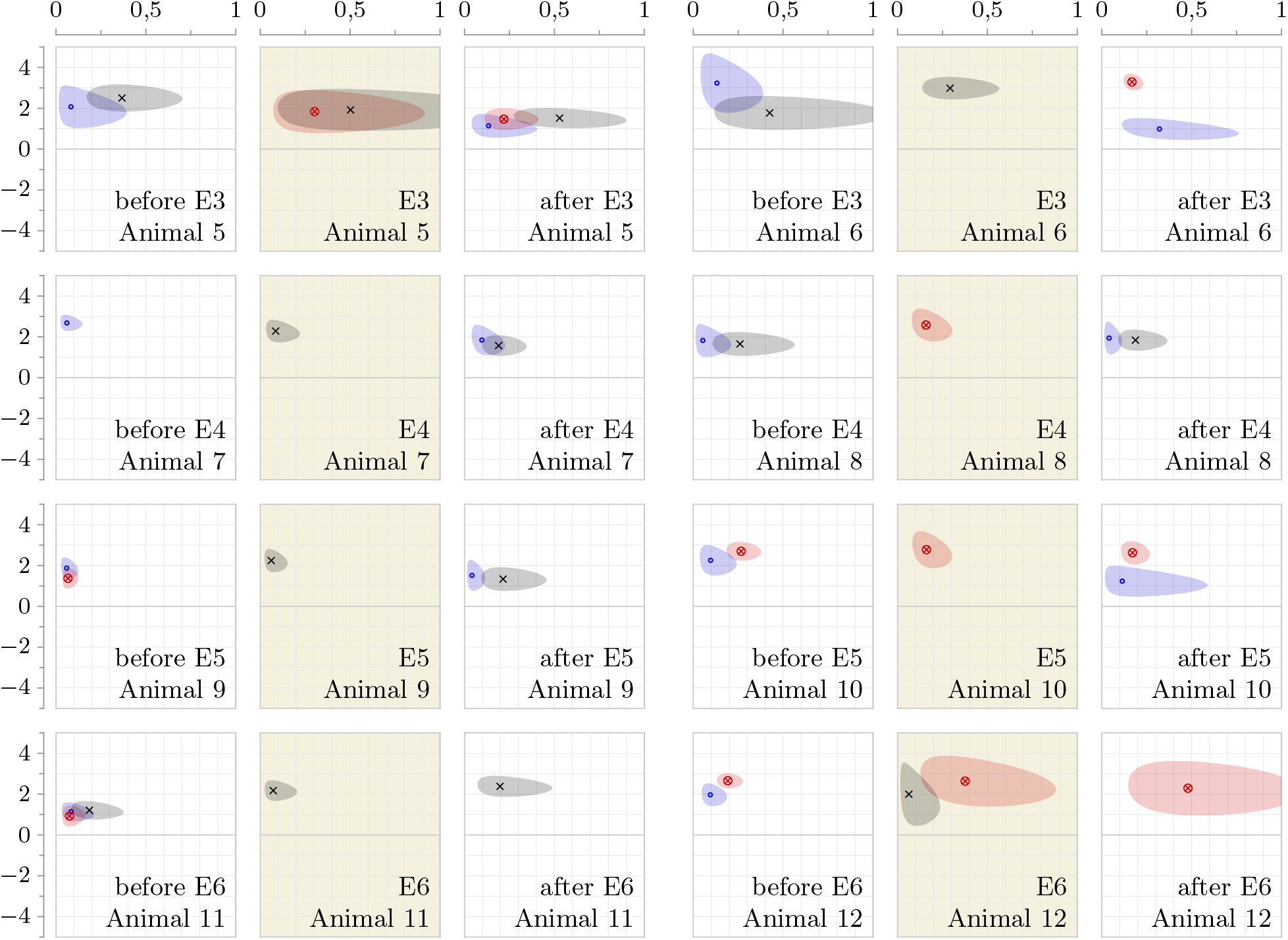
Constellation plots fitted to data subsets covering periods before, during (highlighted) and after minority assignment for eight selected animals.

A clear pattern emerges from the inspection of the plots. All animals start off with a strong social effect, which developed during the adaptation phase A2 in which influence of others was always helpful and continues in the pre-minority period as reflected in model constellations in the leading intervals. Upon change of assignment, mice rapidly change their strategy, relying on reward-based experience in most cases and, in some cases, on the habitual effect. Note that where automodulative locus is present, it comes with an unusually high *α* coordinate. This suggests a strategy consisting in probing a single corner several times, then switching to a new corner. When the minority assignment ends, trust in social influence is restored in most mice. The absence of the social effect in constellations of Animal 11 and Animal 12, for whom the post-minority period covers only one phase, suggests that reinstatement of the social effect requires more time than the duration of the phase.

In conclusion, models of behavior fitted to assignment-relative intervals indicate that mice are able to detect the divergence of the goal of the group and their own and rapidly change their learning strategy in response to this situation.

## 3 Discussion

The importance of social influence in learning has been demonstrated in many earlier studies. For example, in an Intellicage-based experiment, Kiryk et al. (2011) found that while learning was severely impaired in Alzheimer-type transgenic mice, the same mice were able to perform almost as well as the healthy control group when both groups were mixed together [37]. Here, we give an analytical description of social learning within the framework of a plurivalent reinforcement model. Applied to data representing mice behavior in our experiment, the model shows that the decision rules of mice vary depending on majority-minority assignment. When in the majority group, social influence is beneficial and animals readily use it to their advantage, whereas during their minority assignment, the activity of others is largely ignored as misleading. This suggests the existence of a brain mechanism able to evaluate the usefulness of the particular elements of the decision making policy and to adjust their respective strengths. Such an adaptation mechanism can explain the results reported by Kiryk et al. and others [17], and is consistent with the theory of social niche specialization [45].

In our experiment, the observed strategy reflected how focal’s goal related to majority. The relation between own and other’s goals has been the object of previous human studies. Lebreton et al. [19] demonstrated the mimetic desire effect, or the preference for objects that are explicit goals of others. Charpentier et al. [20] considered how hidden goals are indirectly communicated by observed choice and which of a number of policies best fits behavior, depending on relative uncertainty between observation and direct experience. Colette et al. [46] framed imitation as inverse reinforcement learning in a task where the goal of artificial demonstrator was fixed. In machine learning, imitation learning is an approach to which one may resort for problems too difficult to solve in associative way through trial and error, such as playing chess or autonomous driving, but for which demonstration data from human performing the task are available [7]. Its simplest form is unconditional behavior copying, which reduces to a supervised learning problem, but has the disadvantage of reproducing the flaws and weaknesses of the demonstrator’s policy. In contrast, inverse RL attempts to recover an approximation of the latent reward signal from the apparently value-free observed behavior [47] and through this to infer the goal of the observed agent.

By adopting action-value reinforcement model of imitation, we assumed causal and directed influence relations between each pair of individuals forming the group. An earlier study in mice by Smutek et al. [42] mapped pairwise social interaction measured by the rate of visits in a corner directly following a visit by another individual. Similar methods but in a collective setting were used later [48, 29]. While these measures certainly relate to social influence, they fail to reflect its lasting effect, as they materialize only in the animal that first follows the previous choice, disregarding subsequent visits. Other relevant works mapping pairwise social relations across the cohort measured different quantities such as dominance relation [49] or place cooccupation (an undirected relation) [50, 51].

The assumption that mice are able to estimate the current choice preference of every other group member may hold in very small groups but is implausible at greater scale. When the social environment is too complex, animals might restrict their attention to a subset of influential individuals or use the collective variant of social valuation. Our modeling results do not give a decisive indication in favor of either of these possibilities. Some mice seem to be socially selective while others treat the examples from others more homogeneously, thus favoring the collective model of influence, yet the differences in model likelihoods are modest. Variation in the predictive power of observations from different individuals, which we exploit as the ‘exclusion loss’ metric, suggests the possibility of collective valuation dynamics applied to observations filtered on a simple “who” basis. A radical simplification is proposed by Najar et al. [29], namely that the agent maintains a single valuation aggregating both observed actions of others and outcomes of own experience, with different learning rates. While these autors argue that such an aggregation best explained observed behavior, this may stem from the peculiar experiment design in which two demonstrators in a binary choice task in fact had opposite goals, and observation of either could impart the correct choice. In our scheme where animals face four choices, observation of an individual with a mismatched goal is not as informative.

Machine learning distinguishes model-based and model-free approaches depending on whether or not state structure of the environment is explicitly represented in the agent model. Our experiment with immediate reward is designed for model-free description. While there are no environmental (i.e., external) states to be represented in the decision making model, we discover distinct behavioral patterns depending on observed drinking condition, suggesting the existence of two states internal to animals, which we link with the motivation to drink. Thus, animals alternate between the associated decision rules, resulting in complex behavioral patterns which could be disentangled in our study. Existence of other latent states can be predicted, for example related to attention. We model mice as omnivigilent agents, indefatigably tracking others’ corner preference at all times. In reality, mice must rest from time to time. Moreover, they cannot observe others while inside a corner (a measurable circumstance which we take into consideration in our companion paper [38]). It is conceivable that some effects we observe, e.g. the residual negative automodulation in some mice, are linked with attentional states or other animal-internal variables which remain obscured.

We also observe selective imitation of choices of others depending on their drinking condition. This condition and the associated motivational state could be inferred from the durations of visits, longer in general when animals drink, yet we can cannot rule out the possibility of other involuntary cues or deliberate signals. Winiarski et al. [49] reported olfactory signaling of reward in mice, albeit in a different housing condition which could maintain these signals for a longer time. Whether the same mechanism could work within the Intellicage is uncertain due to its small size, while odors are arguably distributed in space.

Midbrain dopaminergic neurons were shown to act as the primary neuronal substrate of reinforcement [52, 53]. More recently, a link to midbrain dopamine has been established in apparently value-free phenomena such as perseveration [44, 13], preference for relative novelty [12, 31] and imitational learning [19], all of which are reflected in our plurivalent model. A general theory of the computational function of the dopamine system was proposed by Bogacz [41]. In his theory, the habitual system yields the prior probability, which, together with probability conditioned on past exprience, forms the maximum a posteriori estimate. Bogacz also predicts the existence of a third system, responsible for the prediction of value target (i.e., level of reward attainable in a given state) which, combined with the other two, sets the problem of choice as a fully bayesian probability computation. This third element can serve as the ground for explaining why expectation of reward develops, a phenomenon which we account for in the reward valuation dynamics, without explicitly modeling as a transient process. Within this framework, social imitation should probably be regarded as a component parallel to the habitual system, assisting in the estimation of the prior probability, but on the other hand, our study reveals cases in which reward valuation plays no reinforcing role, but imitation does. Thus, how social learning fits in brain computations is yet to be understood.

## Acknowledgments

BJ, ML, and DKW acknowledge support by the project POIR.04.04.00-00-14DE/18-00 carried out within the Team-NET programme of the Foundation for Polish Science co-financed by the European Union under the European Regional Development Fund. The authors declare no conflict of interest.

## 4 Methods

### 4.1 Animals

Experiments were performed on 14 female C57BL/6 mice from the colony maintained at the Maj Institute of Pharmacology of the Polish Academy of Sciences in Kraków, Poland. Mice were housed in Plexiglas cages (Type II L, 2-5 animals per cage) with aspen bedding (MIDI LTE E-002, Abedd) and nest-building material, under a 12 h light-dark cycle, with an ambient temperature of 22 ± 2°C. Animals had free access to water and chow (RM1 A (P), Special Diets Services) and were provided with a piece of aspen wood for chewing. Mice were 11 to 15 weeks of age and weighed 19.3 to 22.3 g at the start of the procedure. Mice were implanted with radio frequency identification chips (RFID chips, UNO PICO ID, AnimaLab, Poland) before being introduced to the Intellicage. All experiments were conducted in accordance with the European Union guidelines for the care and use of laboratory animals (2010/63/EU) and were approved by the II Local Bioethics Committee in Kraków (permit LKE 109/2021). All testing was conducted using Intellicages (New Behavior, Switzerland). The Intellicage consists of a base transparent plastic box (55 × 37.5 × 20.5 cm) with a metal cover and custom corner compartments. Each of the cage corners is a small metal chamber that may be fitted with 250 ml bottles, which become available when a plastic cover is lifted to allow access to the nozzle of the bottle. The size of the corner allowed only one animal to enter. Antennas inside the corners detect the RFID tags carried by mice and transmit the number of the tag to the Intellicage Controller software (TSE, Germany). The following events were recorded: presence of an animal in a corner, the crossing of photocell beams placed in the doors leading to the bottles, and lickometer contacts (the animal closing a circuit between the floor grating and the metal dipper of the bottle).

Testing started with adaptation followed by the main procedure with a choice between water and saccharin bottles. The experimental schedule is outlined in Fig 1. After being introduced to the cage mice were allowed free access to all bottles in all corners for 4 days. This was followed by adaptation to the configuration with one door per corner open, which lasted 4 days with a switch of accessible bottles in every corner after 2 days.

Then, when an animal entered a corner the doors would be initially closed, but after 1 s, one of the doors would open providing access to a bottle. The door would close 10 s after licking was detected or when the animal was no longer detected in the corner. In the final adaptation stage, one of the accessible bottles was filled with 0.1% saccharin solution. The position of the saccharin bottle was changed every 24 hours and for 8 days. Altogether, the adaptation to experimental environment lasted 16 days in total in 3 phases.

During the main experimental procedure, animals again had access to 3 water bottles and 1 saccharin solution bottle, and were split into two groups. 12 animals were assigned to the ‘majority’ group and 2 to the ‘minority’ group. The two groups differed in the position of the accessible saccharin bottle, but had access to water in all corners. The assignment of groups changed every 4 days, in a way that every animal was placed in the minority group for one 4-day period.

### 4.2 Formal definition of the model

The complete description of the behavior of a group of animals (more generally, agents), at the level of detail of interest to us, is given by the sequence of tuples (*t*_*n*_, *A*_*n*_,*C*_*n*_, *R*_*n*_*),n* = *1*, …, *N*, denoting, respectively, the time of visit, the visiting animal, the visited corner and visit outcome. Order in time is assumed: *t*_*n*_ *< t*_*n*+1_ for all trials *n*. This is the formulation of a simple marked point process [54], in which visits are each labeled with the categorical variables *A*_*n*_ ∈ 𝒜, *C*_*n*_ ∈ 𝒞, *R*_*n*_ ∈ {0,1}. Here, 𝒜 denotes the set of *K* animals in the cohort (more generally, agents forming the group) and 𝒞 the set of corners (possible actions). Unlike in our companion paper [38], here we ignore the duration of visits.

We model the decision making behavior of a chosen individual *i*, the focal, under the influence of others’s choices. Using the condition *A*_*n*_ = *i* for each individual *i* ∈ 𝒜, we split the sequence of visits into *K* animal-specific subsequences. Let *t*_*i,n*_, *C*_*i,n*_, *R*_*i,n*_ denote the sequence of visit instants, corner choices and outcomes relative to individual *i*, where now *n* = 1, …, *N*_*i*_. Thus, we have a family of point processes for each individual in the group, where each event is marked with the choice and its outcome.

With these definitions in place, the vector-valued encoding for the experienced reward, (**R**_*i,n*_), is

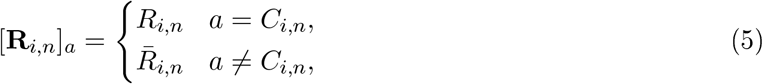

where 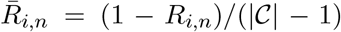 is the fictive reward assumed in the corners not visited. The denominator is chosen to balance the actual and fictive rewards when multiple choices are possible. In a cage with four corners, the denominator value is 3, reflecting the number of possible alternative choices if a wrong action was taken, such that the total fictive reward is 1. The associated valuation is given by the linear first-order recursive dynamics

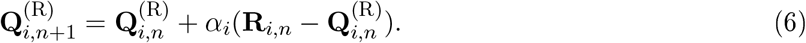

This vector-valued valuation reflects the effect of experienced reward.

We further introduce the social signal (**S**_*j*,*m*_) emitted by animal *j:*

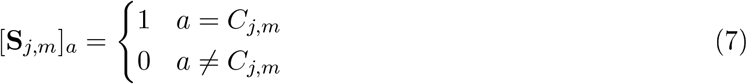

It is the imitational equivalent of Equation (5), where each visit of *j* advertises the chosen corner, *C*_*j*,*m*_. The effect of social influence of *j* on *i* is based on the valuation rule derived from this signal

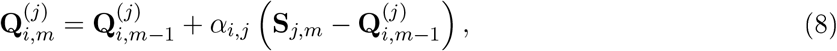

where *α*_*i,j*_ is the learning rate parameter specific to the focal-influencer pair.

The point process structure of the problem is not negligible when effects of different animals are considered: the social signals **S**_*j*,*m*_ and the derived valuations are sampled at time instants different than the focal’s visits. The causality principle requires that choice in a given trial be conditioned only on past events. For this to hold, we define the time alignment as mapping the sequence of valuations 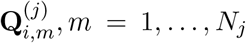 (derived from the social in fluence emitted by animal *j*) into a new sequence 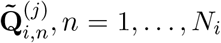 coincident with visits of focal animal *i*, by selecting the last value effective before the time of *i*’s *n*-th visit:

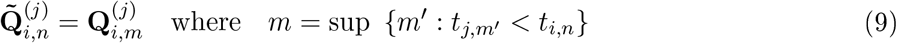

This operation reduces to a one-trial delay in the special case *i* = *j*, reflecting the fact that current choice can be conditioned on past choices but not on itself. In reward-based valuation, the delay is explicitly represented in formula (6) (index *n* + 1 on the left hand side).

Finally, we formulate the model of behavior, or the social reinforcement learning policy, by postulating a linear rule of effects composition

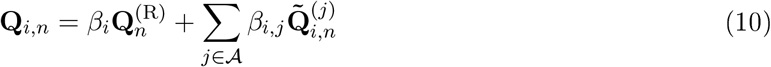

coupled with the conventional logistic map from values to probability

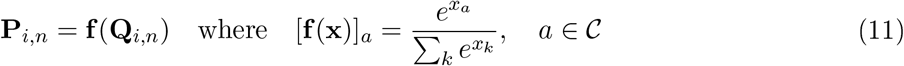

The valuation sum in Equation (10) does not include a constant bias as used in some studies to reflect individual corner preference [16, 35, 30, 18]. Rather than relying on such phenomenological description, the model strives to explain why corner preferences develop through the automodulative term, and why they tend to converge in most animals through social influence.

Depending on the amount of data, one may decide to trim down the model by considering a select subset of the terms in Equation (10). As a special case, retaining only the reward term yields the multiarmed bandit model. Alternatively to selecting a subset of influencers, one may consider the collective influence of the group *G*(*i*) = 𝒜 − {*i*}, obtained from the social signal (**S**_*G*(*i*)*m*_) originating from all animals except the focal taken together. This variant reduces the summation on the right hand side of (10) to just two components, the self effect and the group influence, which yields net valuation of the form

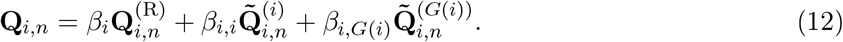

The collective social effect 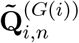 is calculated according to Equation (8) with (**S**_*G*(*i*)*m*_) as input and time-aligned using Equation (9). With three terms and six parameters, this variant obviates the need for effect selection.

### 4.3 Model mechanics

The stochastic prediction rule, Equation (11), relates the probability of corner choice to combined valuation, Equation (10). Values of chosen corners should be high for the model to attain high likelihood. The components of the linear combination, Equation (10), are 4 × *N*_*i*_ matrices, and can be treated as vectors spanning a subspace of the original space, which we can visualize as a projection on a plane (Fig. 9 A).

**Figure 9:**
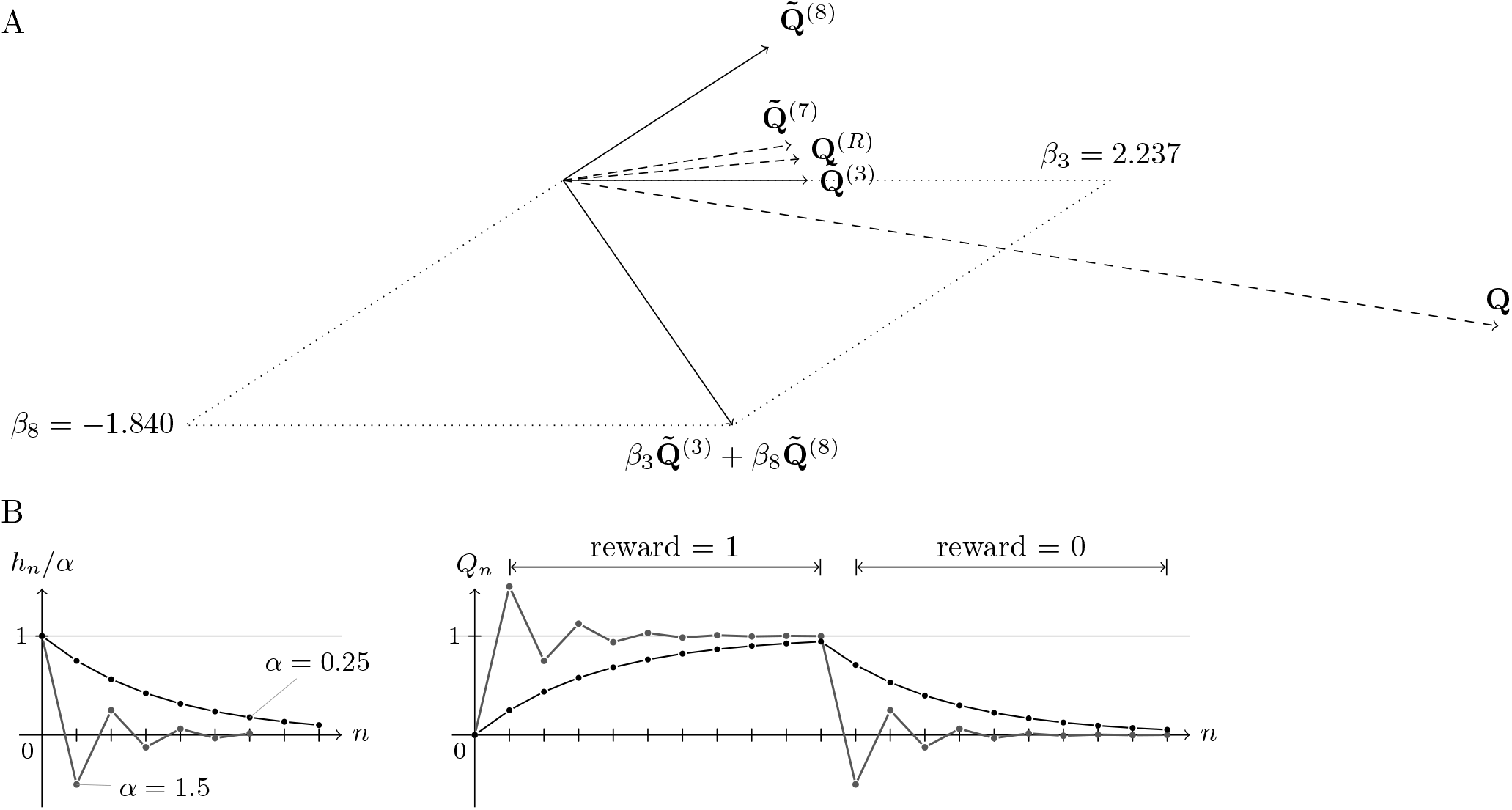
A, Vector mechanics of composition of effects in the social influence model of Animal 3. Solid vectors lie in the plane of the plot, dashed vectors are projections from higher dimensions. B, Valuation impulse response (left) and waveform for piecewise-constant reward input (right) with two chosen learning rates, *α* = 0.25 and *α* = 1.5. The value of the learning rate affects the orientation and magnitude of valuation vectors shown in A.

Let us ignore at first how valuation vectors depend on respective *α* parameters and assume that these parameters are fixed and known. Subject to this simplification, the problem of fitting the plurivalent model reduces to a simple logistic regression on four classes, and amounts to fitting *β* parameters, the strengths of the fixed effects. Under mild conditions which generally hold in sufficiently big samples [55], this problem is convex and has a unique solution. Nevertheless, this solution will not always adequately reflect the decision making process.

Fig. 9 A illustrates an ill-conditioned model in which one of the social effects (influence of Animal 8) has a negative strength (*β* = −1.84). The model was fitted to Animal 3 with no constraints imposed on either *α* or *β* parameters. We observe positive cosine similarity between each pair of effect, and between each effect and the net valuation, depicted as **Q** in the plot. Despite this, *β*_8_, the weight of the effect of Animal 8, is negative. Apparently, this effect has been selected as a model component not because of it being a good predictor; on the contrary, it is one that is the most off the net direction. Negative *β* values in the linear combination, Equation (10), give the possibility of improving the fit by forming a net valuation outside the convex solid angle spanned by the effect vectors included in the model, which often requires considerable positive and negative weights. Thus, an unconstrained model can show biased effects weights, sometimes negative, not reflecting how the effect of interest relates to the decision rule. As a remedy to this problem, we constrain all social effects to have positive *β* parameter value, which can be seen as a form of nonnegative matrix factorization [56], insofar as social effects (and reward) are considered. We do not impose this constraint on the automodulative effect, which may by itself be negative (cf. Fig. 3). In the case of the model depicted in Fig. 9 A, effect selection subject to the above condition yields the same set of effects, but without Animal 8.

When learning rates (*α*) are optimized jointly with weights, both the length and the orientation of component valuation vectors can vary. In general, these vectors follow a curvilinear trajectory as α varies, starting at **Q**^(·)^ = **0** (the assumed initial value) when *α* = 0. This added flexibility contributes nonconvexity and possibly multimodality to the likelihood function.

### 4.4 Fitting procedures

A stochastic model assigns a probability estimate to each sample. Here, it is the probability 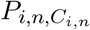 of actions *C*_*i*,*n*_ taken by focal *i* in each trial. The likelihood is the joint probability of the complete sequence of actions, which, by the Markov property, is the product of probabilities in each trial. It is more convenient to express likelihood in the logarithmic scale due to its combinatorial form and to scale it relative to the parameter-free null model assigning uniform probability *P*_*i*,*n*,*a*_ = *P*_null_ = |𝒞|^−1^, which eases interpretation while not changing model estimates

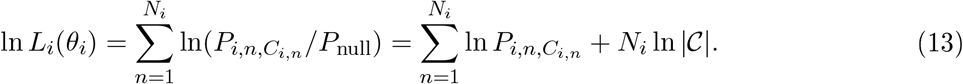

This formula specifies the relative log likelihood (RLL) as a function of the vector *θ*_*i*_ of model parameters, which we encode

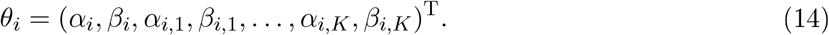

Good models should perform better than the null model and thus attain positive RLL scores. To solve the maximum likelihood estimation (MLE) problem 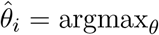 ln *L*_*i*_(*θ*) we resort to gradient optimization, using formulas given in the next section.

Maximizing the likelihood function for all animals *i* ∈ 𝒜 individually yields MLE over the group because models of different animals are conditionally independent given data

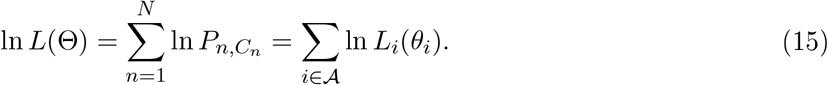

Data collected in Intellicage studies are often intermittent: gaps can occur both due to faults and scheduled actions, such as cage maintenance. Any such gap damages the time series structure of the data and must be adequately handled. We use weighted log-likelihood

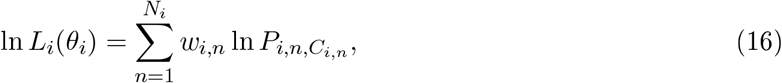

with weights *w*_*i*,*n*_ *=* 1 except during designated gap periods when *w*_*i*,*n*_ *=* 0. To remove likelihood artifacts right after the gap, we let the weights gradually return from 0 to 1 using *w*_*i*,*n*_ *=* sin^2^(*πt*/2*T*_*w*_), where *t = t*_*i*,*n*_ *− t\** is the time elapsed since end of last gap period and we set *T*_*w*_ to 4 hours. We also employ weighting for phase-local fits, simply setting *w*_*i*,*n*_ *=* 0 outside the period of interest (which retains history dependence).

Overfit may occur with insufficient data, often leading to effects with *α* ≈ 0 and an excessive magnitude of *β*, but this can be prevented by regularization of the optimization problem. We use the following objective function including a quadratic penalty on *β* parameters

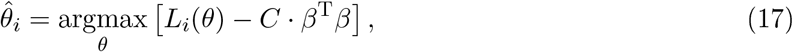

which can be regarded as maximum a posteriori estimation with normal prior on *β*. The meta-parameter *C* controlling the strength of regularization must be set appropriately. There is no theoretical guidance to the choice of this value in general. However, in estimating phase-by-phase models in Fig. 8, we can use the natural reference when models fitted to the entire experimental part (phases E1–E7) are well behaved. First, we calculate *C* incurring a 5% RLL penalty at the fitted parameters. This is an upper bound, since penalized fits have slightly suppressed *β* parameters, although the perturbation of effect loci is minor at this level of regularization. Subsequently, we split the regularization value in proportion to the number of visits in each subset, reflecting the assumption of equal expected RLL contribution of each trial and model stationarity. In reality, constellations of phase-local fits do vary and the corresponding RLL scores are higher than estimated from the proportion.

The likelihood framework lays the foundation for model comparison and selection. Models with many parameters are more flexible and can attain greater likelihood, but are susceptible to overfit. The Bayesian (Schwarz) information criterion (BIC), expresses the goodness of fit adjusted for model size, and is defined as follows:

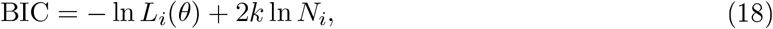

where *k* denotes the number of parameters, *k* = 2(*K* + 1) for a full-order model. Given multiple models, the one with the lowest BIC score is preferred. This rule is applied in conjunction with the following stepwise effect selection procedure:

1. start with an empty model;
2. for each effect not included in the model and for each *α* chosen from a grid on (0, 2): augment the parameter vector by setting *α* and *β* = 0 for the new efect and refit the new candidate model;
3. select the candidate model with greatest likelihood;
4. go to 2. unless all effects are included.

This greedy procedure produces models with increasing number of effects in each step. The candidate model with the lowest BIC is finally chosen.

Besides BIC, we evaluate per-trial accuracy defined as

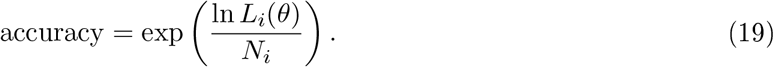

This formula compensates the systematic variation of likelihood with dataset size. It is the geometric average probability of correctly predicting the choice of the focal mouse, considering trial by trial prediction by the agent model as a simple Bernoulli process.

We measure the strength of influence in a pair of animals (from *j* to *i*) using exclusion loss, which is the difference in log-likelihood between two models with collective influence. The reference model has valuation given by Equation (12), where the social effect 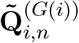 is calculated from the visits of all remaining animals grouped as a single class. The other model has the same valuation formula but with the social effect 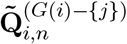 in which visits of j are ignored. Denote the likelihoods of respective models by 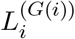 and 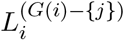. Then the exclusion loss is defined as

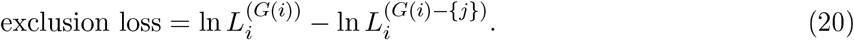

### 4.5 Likelihood: value, gradient and hessian

Let *X*_*n*_ denote one-trial delayed reward or social influence signal, as adequate. Then, with animal and corner indices implied, the valuation formulas for all effects have the general form:

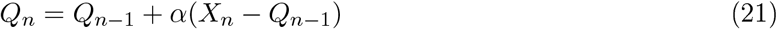

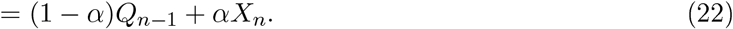

The above expression is a first order autoregressive filter equation with input *X*_*n*_ and output *Q*_*n*_, and can equivalently be written using the convolution summation formula

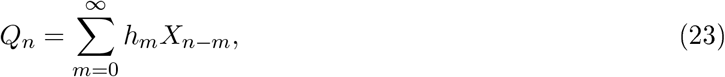

with infinite impulse response (weighting kernel) *h*_*m*_ *= α*(1 − *α*)^*m*^. Stability is assured when *α* ∈ (0, 2). For *α <* 1, the impulse response is a geometrically decaying kernel, leading to monotonic convergence to a limit under constant reward condition (Suppl. Fig. 4.3). In this case, the convergence can be characterized by a time constant *τ*, based on exponential approximation of valuation in continuous time. Using the focal’s average rate of corner visits, *λ*_*i*_ = *N*_*i*_*/T* (ignoring its circadian variation)

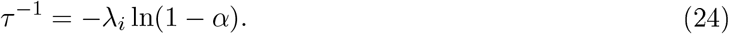

This time constant is a useful diagnostic for *α*: it grows as *α* → 0, but should not exceed the period of reward changes. Excessive time constants are implausible and may signal long-term phenomena unrelated to reinforcement. Thus, *α* should not be too small. The case *α* = 1 makes *h*_*m*_ a unit impulse, and hence *Q*_*n*_ *= X*_*n*_. When *α >* 1, the impulse response decays in an oscillating manner and leads to value overshoots (Suppl. Fig. 4.3) which can be interpreted as overexpectation. The oscillations decay quickly unless *α* considerably exceeds unity, which is again an implausible condition.

Taking the signal processing view of valuation dynamics is useful for the efficient numerical evaluation of likelihood and its derivatives.

#### Log-likelihood gradient

**grad** ln *L*_*i*_ is the vector of partial derivatives

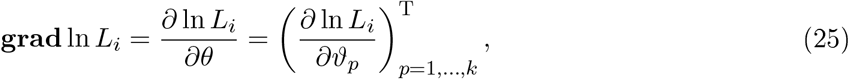

where *ϑ* denotes an arbitrary element of the parameter vector *θ* and

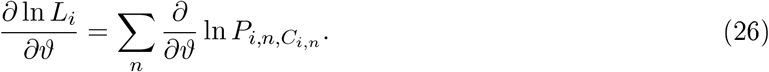

The following holds for all *α* ∈ 𝒞

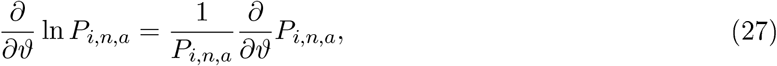

and, from the logistic formula, Equation (11) for **P**_*i,n*_,

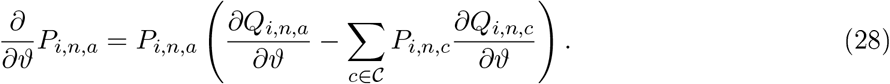

Substituting Equation (28) in Equation (27) yields the general form

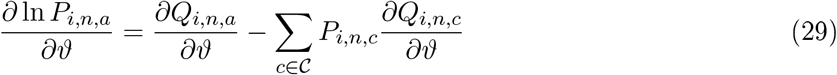

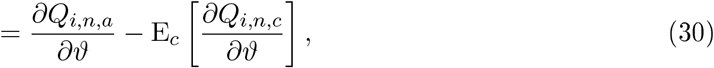

where E_*c*_ denotes expectation over corners. This formula relates the likelihood gradient to the partial derivative of valuation, which remains to be determined separately for each of the model parameters substituted for *ϑ*.

First, consider the effect of reward and its two parameters *α*_*i*_, *β*_*i*_. Taking the partial derivative of the composition formula Equation (10) with respect to *β*_*i*_ yields

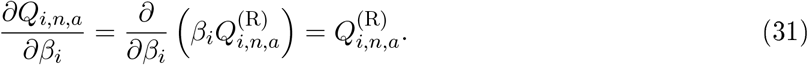

This result combined with Equation (30) gives the solution

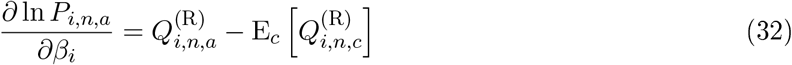

For *α*_*i*_, from the composition formula, Equation (10),

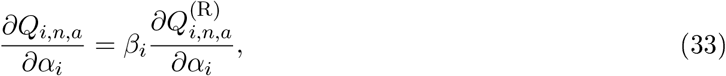

and it follows by differentiation of the update dynamics formula, Equation (6) that

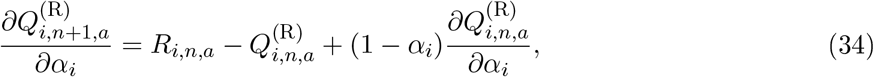

which is a first order autoregressive filter equation with input 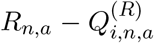 and output 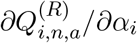, shifted by one trial. Thus, we evaluate 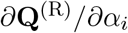 through a filtering operation and feed the result into Equation (30) to obtain *∂* In 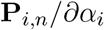.

Calculation of automodulation derivative proceeds analogously, with **R**_*i,n*_ replaced by **S**_*i,n*_. To obtain derivatives of time-aligned social valuations, we first compute the derivative of the unaligned valuation, replacing **R**_*i,n*_ with **S**_*j,m*_, and subsequently apply time alignment, Equation (9), instead of the one-trial shift. This change of order of time alignment and filtering is allowed as both are linear operations.

#### Hessian of the log likelihood

**H** is the matrix of second partial derivatives

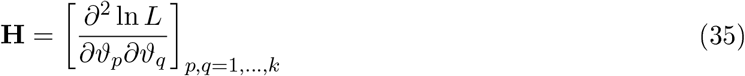

The per-trial contributions to Equation (35) have the general form

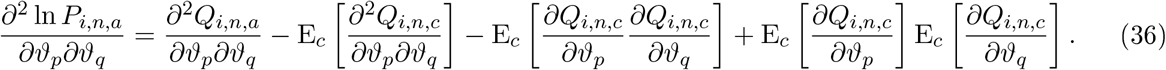

involving second derivatives of *Q* in the first two terms, in addition to the first derivatives, the formulas for which were given. As *Q* is a linear combination of component effects, second derivatives vanish when parameters *ϑ*_*p*_, *ϑ*_*q*_ correspond to two distinct effects. Second order derivatives must be considered in hessian elements corresponding to the same effect, which form 2 × 2 blocks along the diagonal, where three cases are distinguished. Consider again the effect of reward:

1. *∂*^2^ In 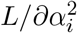: Equation (36) takes the form

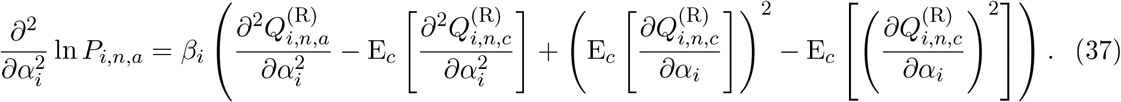 By differentiating Equation (34), we get the identity

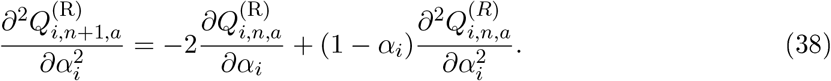 This first order autoregressive filter equation allows to evaluate the second derivative signal 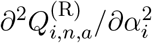 from the first derivative 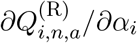. Both quantities fed to Equation (37) yield the desired second derivative with respect to *α*_*i*_.
2. *∂*^2^ In *L/∂α*_*i*_*∂β*_*i*_: by differentiating Equation (32), the partial derivative reduces to

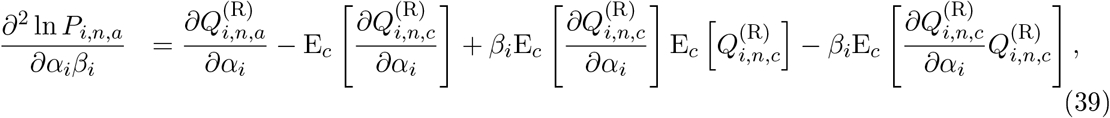

where we again used Equation (28). Thus, only valuation and its first derivative remain.
3. *∂*^2^ In 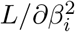: the second derivatives in Equation (36) vanish, leaving only the first derivative in the formula

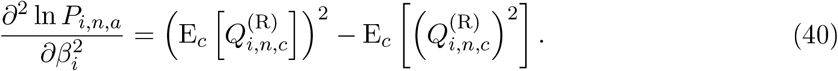

These formulas specify the second derivatives of the likelihood function with respect to the parameters *α*_*i*_, *β*_*i*_ corresponding to the effect of reward **Q**^(R)^. Analogous calculations apply to the remaining effects **Q**^(*j*)^, where **S**_*j,m*_ must be used in place of **R**_*j,n*_. Time alignment operator, Equation (9), is additionally required for social effects.

### 4.6 Confidence regions and statistical significance

Parameter variance is estimated using normal appoximation of the likelihood around a maximum. The covariance matrix is related to the hessian as **Σ** = − **H**^−1^, where we use the sample estimate of **H** at 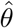 given by the formulas in the previous section. The covariance matrix defines ellipsoidal confidence regions around the maximum likelihood estimate in the parameter space, the boundary of which is the set of points fulfilling

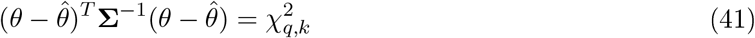

*k* denotes the number of model parameters and the quantile order *q* sets the confidence level; depending on its value and the loci of estimated effects, such regions may extend outside of the valid range of *α* parameters. This makes it necessary to apply a variable transformation. To account for the skew that the likelihood function exhibits in the low *α* region, we relate each a to an auxiliary variable *γ* ∈ ℝ using the logistic function

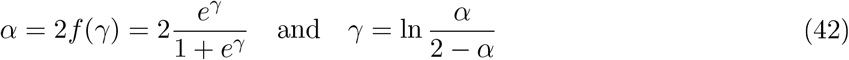

The derivatives *∂F/∂α* and *∂*^2^*F/∂α*^2^, where *F* denotes the (possibly weighted) log-likelihood or its penalized counterpart, Equation (17), require a correction to reflect the transformed parameter domain. The necessary derivative adjustments are based on the identity

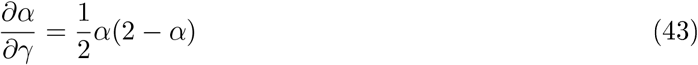

Confidence regions determined in the transformed space are finally mapped back to the original domain using (42) for each *α*.

In practice, only one- or two-dimensional marginal confidence regions (intervals or ellipses) are of interest. Two-dimensional confidence regions which we visualize around effect loci are obtained from marginal covariances extracted as 2 × 2 diagonal blocks (assuming (14) as parameter vector encoding) and the appropriate quantile 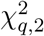. Confidence intervals can easily be obtained from the diagonal elements of the sample covariance matrix (i.e, the variances), from which one calculates interval endpoints by an affine transformation of appropriate quantiles of the standard normal. The dual problem of significance testing of model effects can be based on null hypothesis *H*_0_: *β* = 0 and reduced to interval estimation. Note that, by virtue of the transformation (42), *α* will always be nonzero and cannot be used for testing effect sigificance.

## 5 Supplementary materials

**Figure 10:**
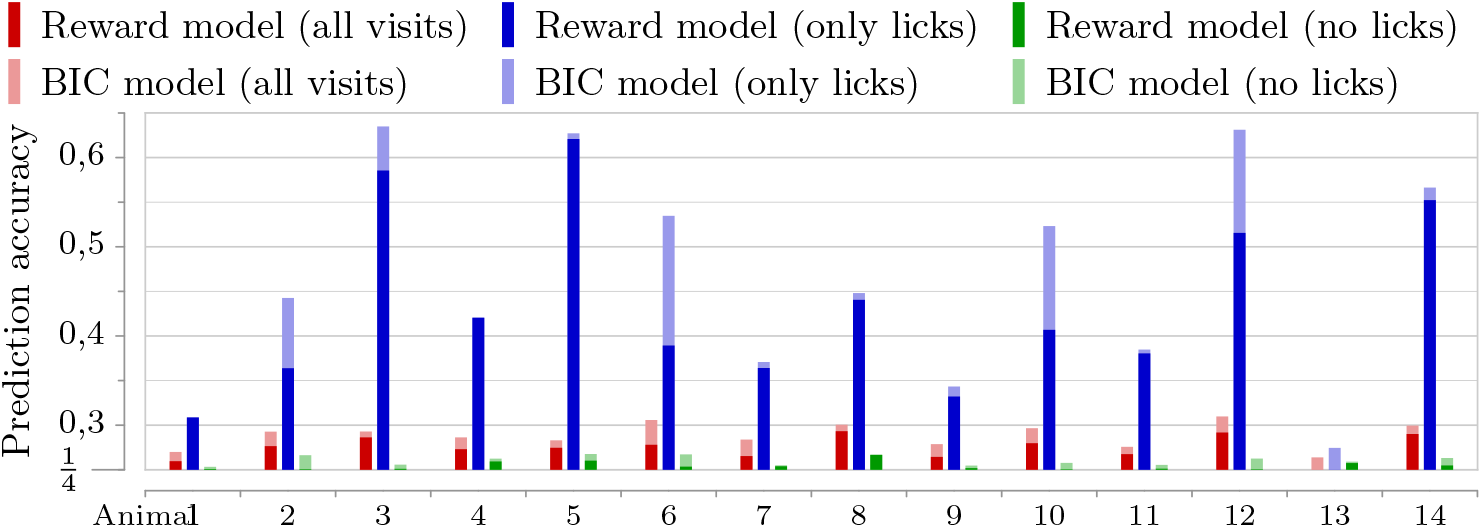
Reclassification accuracy of reward and selected multi-effect models fitted to the full set of visits and its two subsets defined by lick condition.

**Figure 11:**
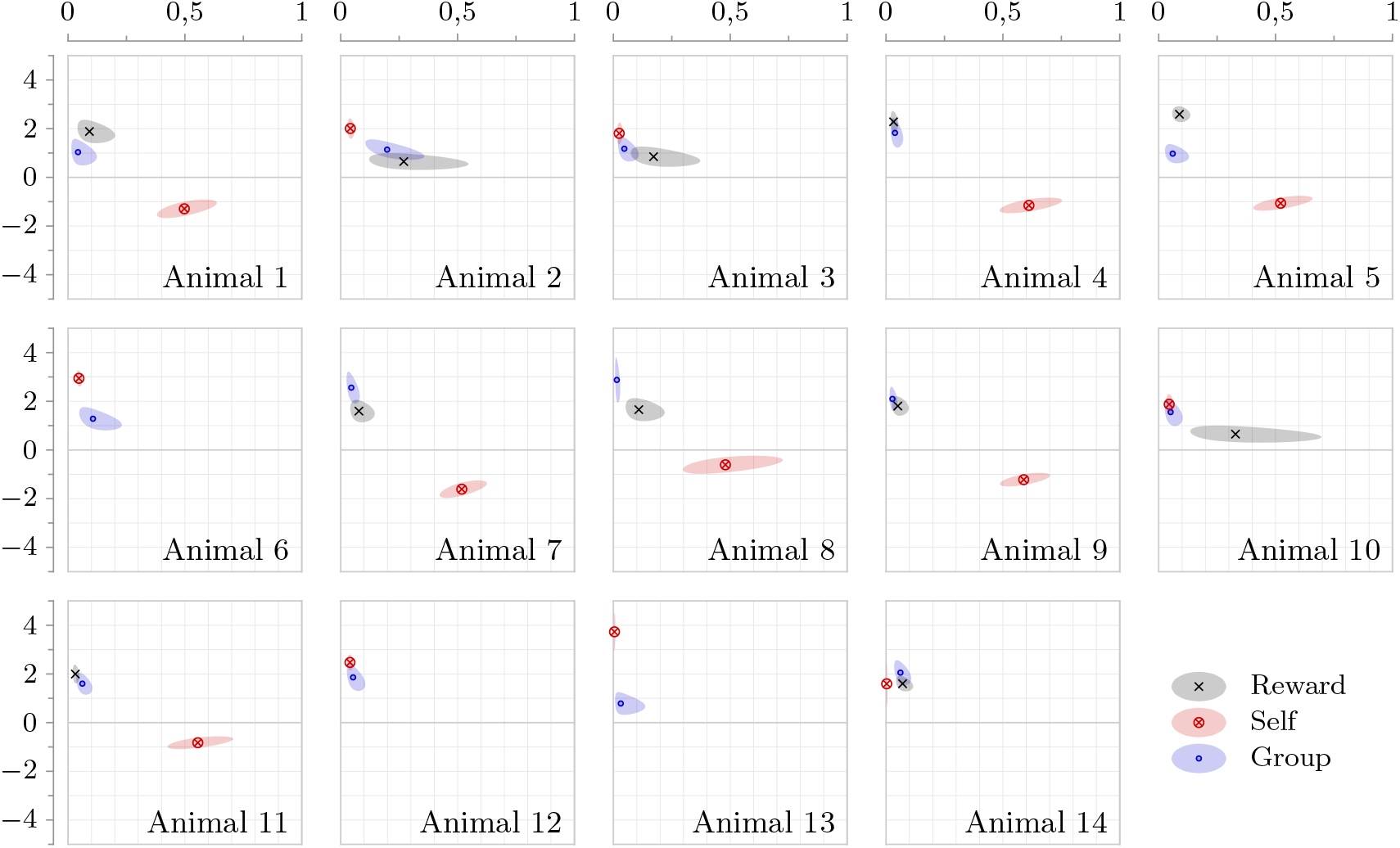
Constellation plots representing models of individual mice, fitted to the set of all visits in phases E1–E7.

**Figure 12:**
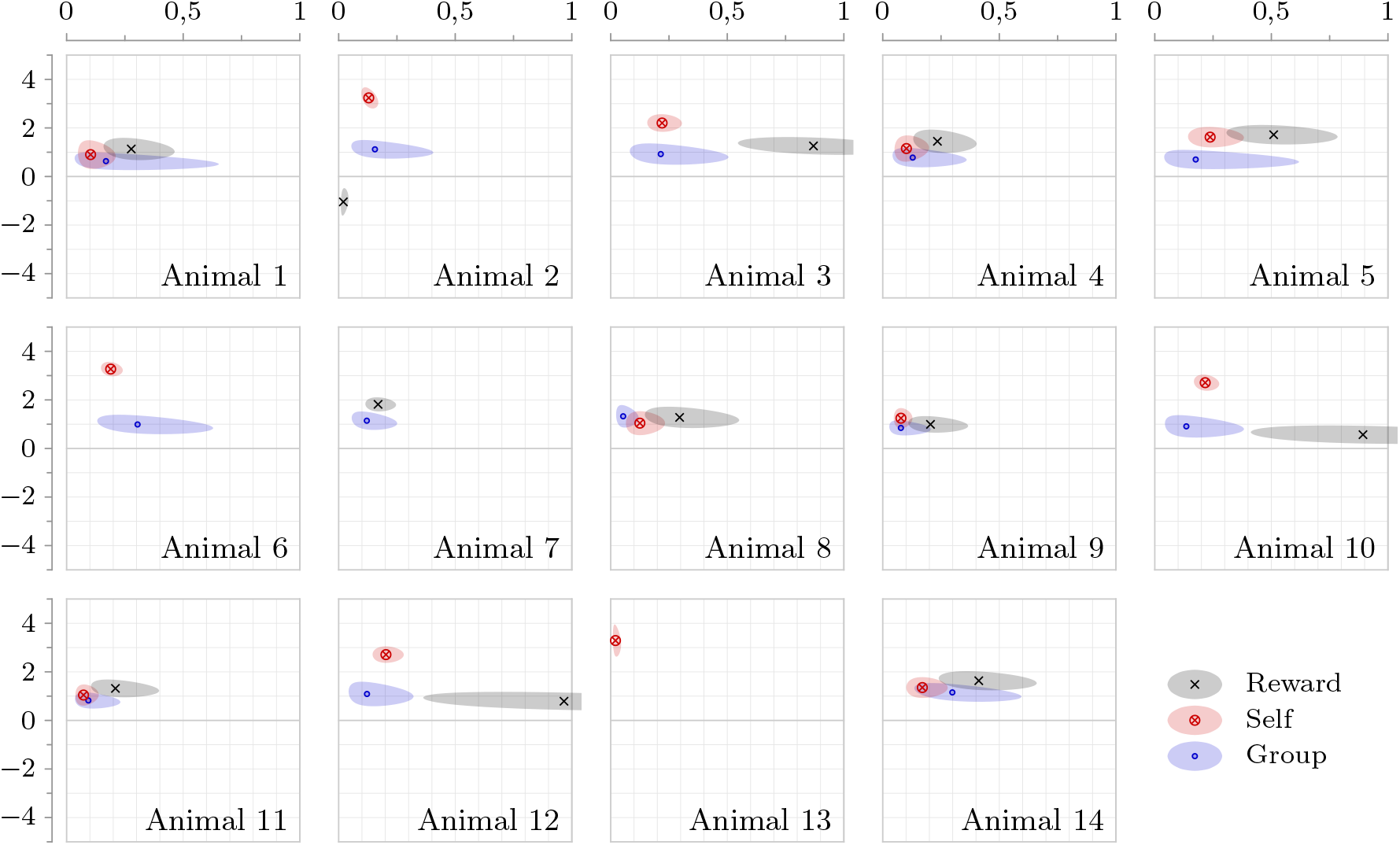
Constellation plots of 5% regularized models of all mice in the cohort, fitted to the subset of visits with detected drinking (goal-oriented condition) in phases E1–E7.

**Figure 13:**
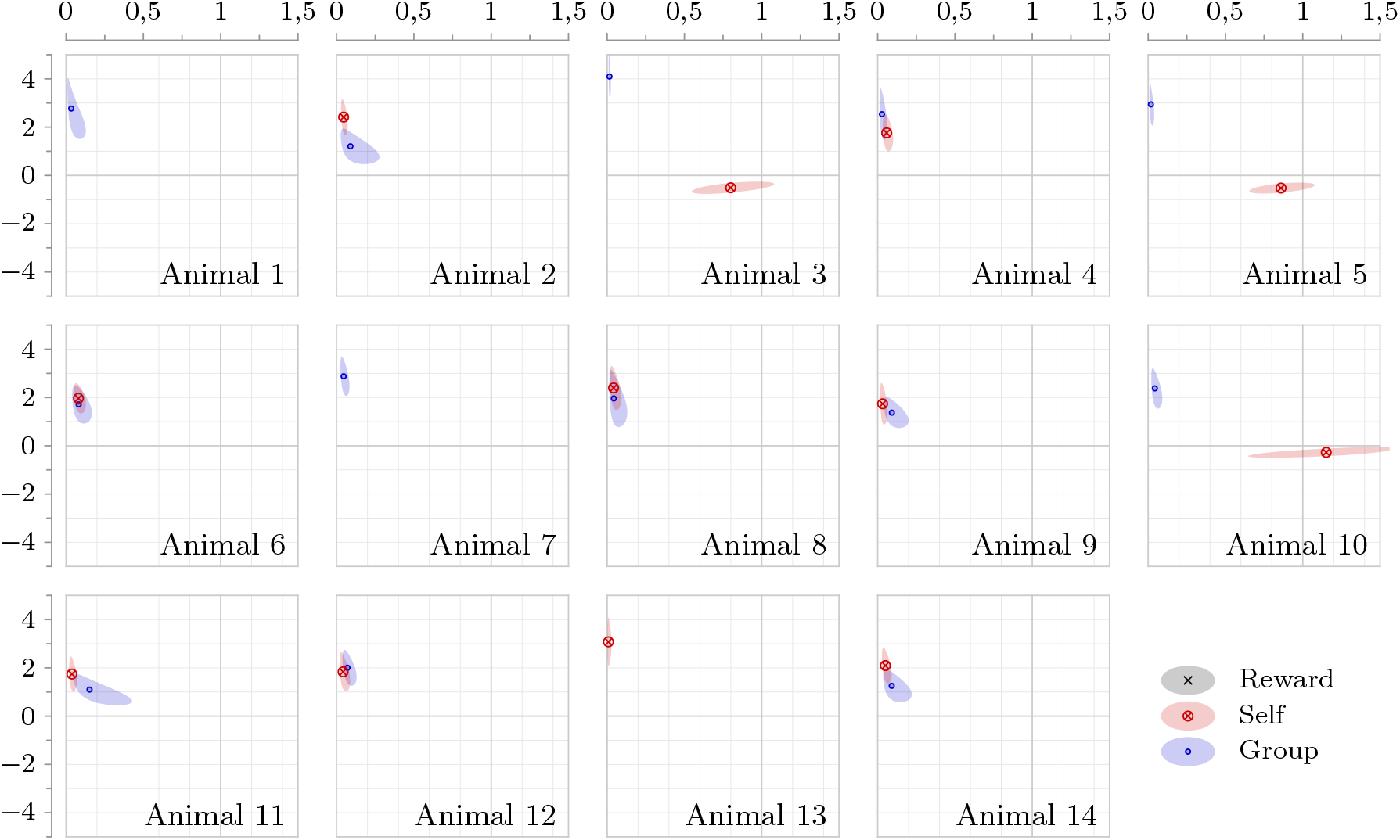
Constellation plots of 5% regularized models of all mice in the cohort, fitted to the subset of visits without drinking (idle condition) in phases E1–E7.

**Table 1:** Visit proportions in corners of the Intellicage by animal and drinking condition.

| Corner | Idle |  |  |  | Goal |  |  |  |
| --- | --- | --- | --- | --- | --- | --- | --- | --- |
|  | 1 | 2 | 3 | 4 | 1 | 2 | 3 | 4 |
| Animal 1 | 21.9 | 25.7 | 24.1 | 28.3 | 28.3 | 24.4 | 27.0 | 20.4 |
| Animal 2 | 23.0 | 28.6 | 22.7 | 25.7 | 23.6 | 36.0 | 24.0 | 16.4 |
| Animal 3 | 20.4 | 28.1 | 25.9 | 25.7 | 19.2 | 40.5 | 12.6 | 27.7 |
| Animal 4 | 22.5 | 22.8 | 27.5 | 27.2 | 15.5 | 37.6 | 18.4 | 28.4 |
| Animal 5 | 24.7 | 25.0 | 22.6 | 27.7 | 30.2 | 20.1 | 33.8 | 16.0 |
| Animal 6 | 26.8 | 25.0 | 23.4 | 24.8 | 42.2 | 32.9 | 18.1 | 6.82 |
| Animal 7 | 26.2 | 26.6 | 25.4 | 21.8 | 18.1 | 32.5 | 19.4 | 30.0 |
| Animal 8 | 24.3 | 19.8 | 35.1 | 20.8 | 17.1 | 36.2 | 19.3 | 27.4 |
| Animal 9 | 20.3 | 25.7 | 27.4 | 26.6 | 22.7 | 26.9 | 27.2 | 23.1 |
| Animal 10 | 20.9 | 29.4 | 24.3 | 25.3 | 20.6 | 35.8 | 17.3 | 26.4 |
| Animal 11 | 30.0 | 24.7 | 21.9 | 23.4 | 17.6 | 28.9 | 19.5 | 33.9 |
| Animal 12 | 18.7 | 29.4 | 23.3 | 28.5 | 12.6 | 37.5 | 11.9 | 38.0 |
| Animal 13 | 15.4 | 33.3 | 24.7 | 26.6 | 16.6 | 26.8 | 17.9 | 38.8 |
| Animal 14 | 18.6 | 28.6 | 23.0 | 29.8 | 23.8 | 25.2 | 30.8 | 20.2 |
| Average | 22.4 | 26.6 | 25.1 | 25.9 | 22.0 | 31.5 | 21.2 | 25.2 |

## Notes

### Competing Interest Statement

The authors have declared no competing interest.

